# Direct interaction between miR-210-5p and HIF-1α regulates HIF-dependent transcription

**DOI:** 10.64898/2026.08.04.742706

**Authors:** Xavier Blanchet, André Mourão, Bartolo Ferraro, Leonardo Matta, Henrika Jodeleit, Phillip von Hundelhausen, Jan Caca, Michael Sattler, Alexander Bartelt, Christian Ries, Christian Weber, Virginia Egea

## Abstract

Hypoxia-inducible factors (HIFs) coordinate cellular adaptation to oxygen deprivation, yet whether hypoxia-induced microRNAs directly regulate HIF-dependent transcription remains unknown. Here, we identify miR-210-5p as a nuclear hypoxamiR that directly binds HIF-1α and enhances HIF-dependent transcription. Hypoxia induced rapid, HIF-dependent nuclear accumulation of mature miR-210-5p across multiple cell types. Biophysical analyses demonstrated direct interaction between miR-210-5p and the HIF-1α bHLH–PAS domain, while mutational studies identified a conserved 5′motif required for HIF-1α binding but dispensable for repression of the canonical cytoplasmic target ISCU. Functionally, transcriptional activity closely correlated with HIF-1α-binding affinity, as binding-deficient variants failed to activate HIF reporters or endogenous target genes. Although AGO2 contributed to hypoxic transcriptional responses, miR-210-5p interacted directly with HIF-1α independently of AGO2. In ischemic myocardium, nuclear enrichment of miR-210-5p and its proximity to HIF-1α support the physiological relevance of this mechanism, revealing a previously unrecognized RNA-mediated layer of HIF transcriptional regulation.

## Introduction

Cellular adaptation to reduced oxygen availability is orchestrated by hypoxia-inducible factors (HIFs)^1^, transcription factors that regulate metabolic remodeling, angiogenesis, and survival pathways across diverse tissues. Although stabilization of HIF-1α is a conserved hallmark of the hypoxic response, HIF-dependent transcription varies substantially between cell types and physiological contexts.^2–4^ This variability suggests the existence of nuclear regulatory mechanisms that fine-tune transcriptional output independently of HIF-1α stabilization.

Given that hypoxic signaling induces a characteristic set of microRNAs (miRNAs) across diverse cellular contexts, we investigated whether hypoxamiRs might participate in nuclear regulation of HIF-dependent transcription.^5,6^

miRNAs are well-established modulators of hypoxic adaptation. Among them, miR-210^7^ is the most consistently induced hypoxamiR and regulates mitochondrial adaptation through canonical post-transcriptional repression of iron–sulfur cluster assembly enzyme (ISCU), succinate dehydrogenase complex subunit D (SDHD), and other components of oxidative metabolism.^7–10^ Beyond these classical cytoplasmic functions, accumulating evidence indicates that selected miRNAs can localize to the nucleus, where they have been implicated in transcriptional regulation through interactions with promoters, chromatin-associated factors, and transcriptional complexes.^11–14^ Recent studies have expanded the repertoire of nuclear miRNA functions to include promoter-associated transcriptional regulation and chromatin-associated gene control; however, the molecular mechanisms governing these non-canonical activities remain incompletely understood.^12^

RNA structure has emerged as an important determinant of protein recognition and interaction specificity, yet direct mechanistic evidence linking structured miRNAs to modulation of transcriptional regulatory proteins remains limited.^15–17^ Understanding how nuclear miRNAs achieve molecular specificity may therefore reveal previously unrecognized mechanisms of transcriptional regulation.

Here, we identify a non-canonical nuclear function of miR-210-5p in the regulation of hypoxic transcriptional responses. We show that hypoxia drives rapid and HIF-dependent accumulation of mature miR-210-5p in the nucleus across multiple cell types. Using complementary biophysical and functional approaches, we provide evidence that miR-210-5p directly engages the HIF-1α basic helix–loop–helix Per–ARNT–Sim (bHLH–PAS) domain^18^ through a conserved 5′ interaction motif. Disruption of this motif selectively impairs HIF-1α binding and abolishes transcriptional potentiation while preserving canonical cytoplasmic target repression, functionally uncoupling nuclear and post-transcriptional activities.

These findings support a model in which direct RNA–protein interaction is required for transcriptional modulation and reveal a non-linear relationship between interaction strength and transcriptional output. Together, our results identify miR-210-5p as a nuclear hypoxamiR that directly engages HIF-1α through a conserved 5′ motif-dependent interaction and establish a mechanism by which selected nuclear miRNAs can modulate transcriptional output through association with regulatory proteins.

## Results

### Hypoxia induces rapid nuclear accumulation of miR-210-5p

To investigate whether hypoxamiRs contribute to nuclear regulation of HIF-dependent transcription, we analyzed the subcellular distribution of mature miRNAs during hypoxic signaling. Pharmacological HIF stabilization with IOX-2 induced marked nuclear enrichment of mature miR-210-5p, whereas miR-126-5p and let-7f remained predominantly cytoplasmic or showed no comparable nuclear accumulation (Fig. 1a). Fraction purity was confirmed by enrichment of Lamin B1 in nuclear fractions and Tubulin in cytoplasmic fractions (Supplementary Fig. 1a).

**Figure 1.**
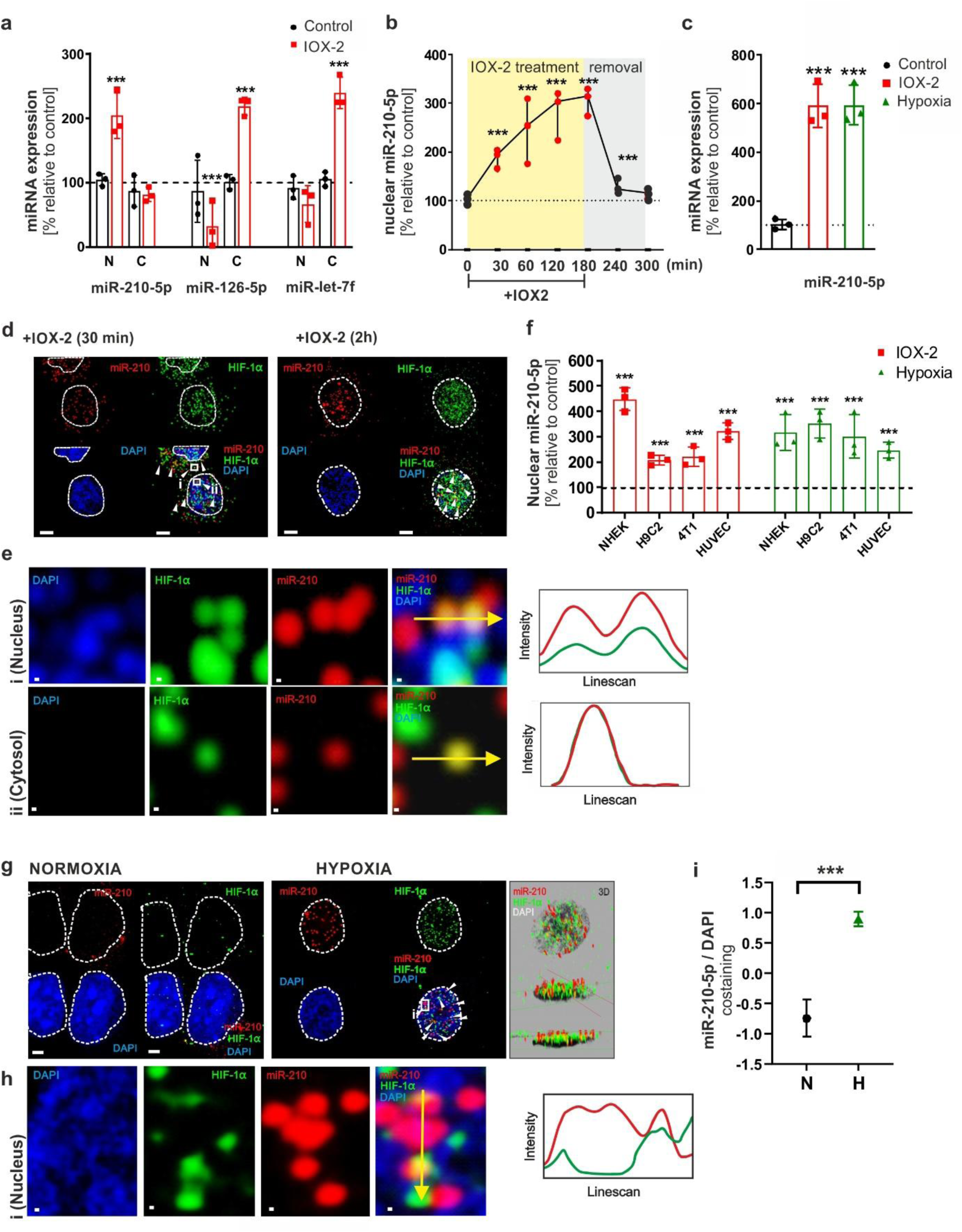
Hypoxia induces nuclear accumulation of miR-210-5p and alters its spatial distribution relative to HIF-1α. (a) Subcellular distribution of hypoxamiRs following HIF stabilization. Primary human keratinocytes were treated with IOX-2 (50 µM, 2 h), followed by nuclear and cytoplasmic fractionation. Relative levels of miR-210-5p, miR-126-5p, and let-7f were quantified by qPCR and normalized to U6. (b) Time-dependent analysis of nuclear miR-210-5p levels following transient HIF stabilization. Keratinocytes were treated with IOX-2 (50 µM) for the indicated time periods followed by compound removal. Nuclear miR-210-5p levels were quantified by qPCR and expressed relative to untreated controls. (c) Analysis of nuclear miR-210-5p abundance under pharmacological and physiological hypoxia. Keratinocytes were treated with IOX-2 (50 µM, 2 h) or exposed to hypoxia (1% O₂, 24 h), followed by nuclear RNA isolation and qPCR analysis. (d) Representative confocal microscopy images of keratinocytes treated with IOX-2 (50 µM) for 30 min or 2 h. miR-210-5p was detected by RNA-FISH (red), HIF-1α by immunofluorescence staining (green), and nuclei by DAPI staining (blue).(e) Representative fluorescence intensity profiles of miR-210-5p and HIF-1á across nuclear (i) and cytosolic (ii) compartments following IOX-2 treatment. Fluorescence intensity profiles were measured along the yellow line. (f) Nuclear accumulation of miR-210-5p under pharmacological and physiological hypoxia across multiple cell types. NHEK, H9C2, 4T1, and HUVEC cells were treated with IOX-2 (50 µM, 2 h) or exposed to hypoxia (1% O₂, 24 h), followed by nuclear RNA isolation and qPCR analysis of miR-210-5p abundance. (g) High-resolution fluorescence intensity profiles of miR-210-5p (red) and HIF-1á (green) under normoxic or hypoxic conditions (1% O₂, 24 h) with orthogonal views and three-dimensional reconstructions. (h) High-resolution fluorescence intensity profiles of miR-210-5p and HIF-1á within nuclear regions under hypoxic conditions. Fluorescence intensity profiles were measured along the yellow line. (i) Quantification of Pearson’s correlation coefficients between miR-210-5p and HIF-1α fluorescence signals under normoxic (N) and hypoxic (H) conditions. Data are presented as mean ± s.e.m. Statistical significance was determined using unpaired two-tailed Student’s t-test or one-way ANOVA with appropriate multiple-comparisons correction. Scale bars, 10 µm unless otherwise indicated.

Nuclear accumulation of miR-210-5p occurred rapidly following IOX-2 treatment and increased over time, reaching maximal levels during sustained HIF stabilization. Removal of IOX-2 reversed this effect, indicating that nuclear enrichment of miR-210-5p is dynamic and coupled to hypoxic signaling rather than reflecting irreversible redistribution (Fig. 1b). Similar nuclear accumulation was observed under physiological hypoxia (1% O₂), supporting the relevance of this response beyond pharmacological HIF stabilization (Fig. 1c).

RNA-FISH and confocal microscopy confirmed spatial redistribution of miR-210-5p during HIF activation. Early after IOX-2 treatment (30 minutes), miR-210-5p signals were already detectable within nuclear regions, whereas HIF-1α remained partly cytoplasmic. Prolonged IOX-2 exposure (2 hours) induced prominent nuclear accumulation of HIF-1α together with increased spatial proximity between HIF-1α and nuclear miR-210-5p puncta (Fig. 1d). Representative fluorescence intensity line-scan analyses demonstrated overlapping nuclear fluorescence profiles of miR-210-5p and HIF-1α, whereas cytosolic regions showed less coordinated signal distribution (Fig. 1e). The nuclear enrichment response was conserved across multiple cellular contexts. IOX-2 treatment and physiological hypoxia increased nuclear miR-210-5p levels in primary keratinocytes, H9C2 cardiomyoblasts, 4T1 breast cancer cells, and HUVECs (Fig. 1f). Under physiological hypoxia, orthogonal views and three-dimensional reconstructions demonstrated focal nuclear accumulation of miR-210-5p in close proximity to HIF-1α (Fig. 1g). High-resolution fluorescence profiles further supported overlapping nuclear signal distribution between miR-210-5p and HIF-1α (Fig. 1h). Quantitative colocalization analyses confirmed increased nuclear miR-210-5p/HIF-1α spatial association under hypoxic conditions (Fig. 1i). These findings identify miR-210-5p as a dynamically regulated nuclear hypoxamiR that accumulates during HIF activation, providing the basis for investigating a potential functional interaction with HIF-1α.

### miR-210-5p directly associates with HIF-1α

Given the pronounced nuclear accumulation of miR-210-5p during HIF activation, we next investigated whether miR-210-5p directly interacts with HIF-1α. HIF-1α knockdown markedly reduced nuclear miR-210-5p enrichment following IOX-2 treatment, indicating that HIF-1α contributes to nuclear accumulation of miR-210-5p (Fig. 2a). Likewise, pharmacological inhibition of importin α/β-mediated nuclear transport with ivermectin significantly reduced nuclear accumulation of miR-210-5p, consistent with importin-dependent nuclear transport contributing to its hypoxia-induced redistribution. Efficient HIF-1α depletion was confirmed by immunoblot analysis (Supplementary Fig. 2a).

**Figure 2.**
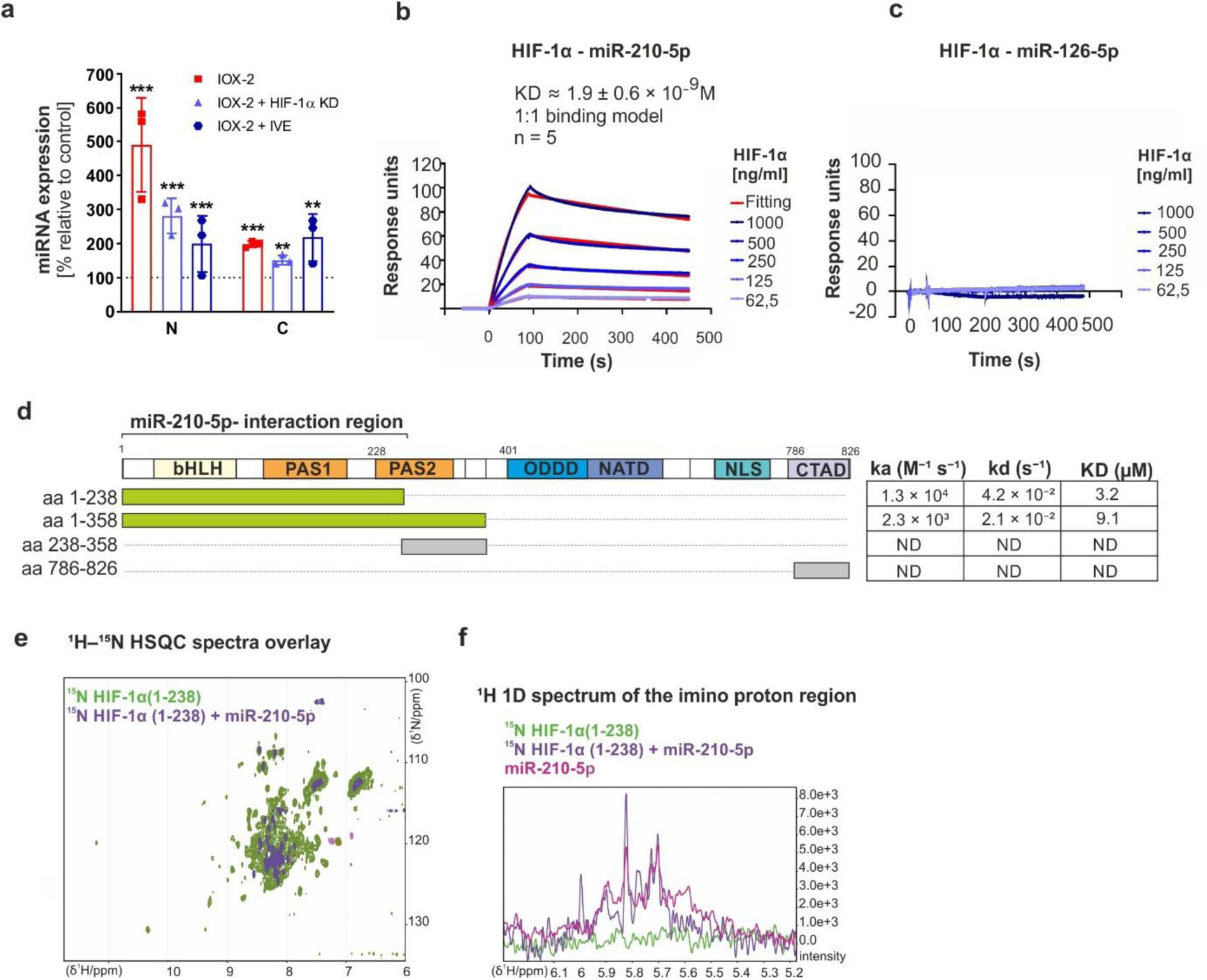
HIF-dependent nuclear accumulation and direct interaction of miR-210-5p with HIF-1α. (a) HIF-dependent nuclear accumulation of miR-210-5p. Primary human keratinocytes were treated with IOX-2 (50 μM, 2 h) alone or in combination with HIF-1α knockdown (HIF-1α KD) or ivermectin treatment (IVE), an inhibitor of importin α/β-mediated nuclear transport. Nuclear (N) and cytoplasmic (C) RNA fractions were isolated and analyzed by qPCR for miR-210-5p expression relative to untreated control (b). Surface plasmon resonance (SPR) analysis demonstrating direct interaction between recombinant HIF-1α and miR-210-5p. Representative sensorgrams and fitted curves based on a 1:1 binding model are shown. Increasing concentrations of recombinant HIF-1α were injected as indicated. (c) SPR analysis of HIF-1α interaction with control miR-126-5p. Representative sensorgrams showing absence of detectable binding between recombinant HIF-1α and miR-126-5p under identical experimental conditions. (d) Schematic representation of the HIF-1α domain organization and recombinant truncation constructs used for SPR binding analyses with miR-210-5p. Summary of ka, kd, and KD values derived from fitting SPR sensorgrams to a 1:1 Langmuir binding model. ND, no detectable binding. (e, f) Overlay of ^1H–^15N HSQC spectra and imino proton spectra of isotopically labeled HIF-1α(1–238) recorded in the absence and presence of miR-210-5p, showing changes in resonance positions and intensities following RNA addition. Data are presented as mean ± s.e.m. from independent biological replicates. Statistical significance was determined using one-way ANOVA with appropriate multiple-comparisons correction.

Surface plasmon resonance (SPR) analysis demonstrated direct interaction between recombinant HIF-1α and mature miR-210-5p with nanomolar affinity (Fig. 2b), whereas the unrelated control miRNA miR-126-5p showed no detectable interaction (Fig. 2c). To define the interaction interface, we analyzed a series of recombinant HIF-1α truncation constructs. SPR mapping localized the interaction predominantly to the N-terminal region encompassing amino acids 1–238 and 1–358, whereas more distal regions showed no detectable binding to miR-210-5p (Fig. 2d). Nuclear magnetic resonance (NMR) spectroscopy provided independent evidence for interaction between miR-210-5p and the HIF-1α bHLH–PAS domain. Addition of miR-210-5p induced signal broadening and resonance loss, consistent with RNA–protein complex formation (Fig. 2e, f).

Collectively, these complementary biophysical approaches demonstrate that miR-210-5p directly engages the N-terminal bHLH–PAS region of HIF-1α.

### miR-210-5p 5′ motif potentiates HIF-1α activity

To define the molecular determinants underlying the interaction between miR-210-5p and HIF-1α, we generated a series of mutant miR-210-5p constructs carrying progressive nucleotide substitutions within the conserved 5′ region (Fig. 3a). RNA secondary structure predictions generated using RNAfold showed progressive alteration of the predicted 5′ interaction element across the mutant series (Supplementary Fig. 3a). The introduced substitutions were designed to perturb this region while maintaining the overall miRNA sequence framework.

**Figure 3.**
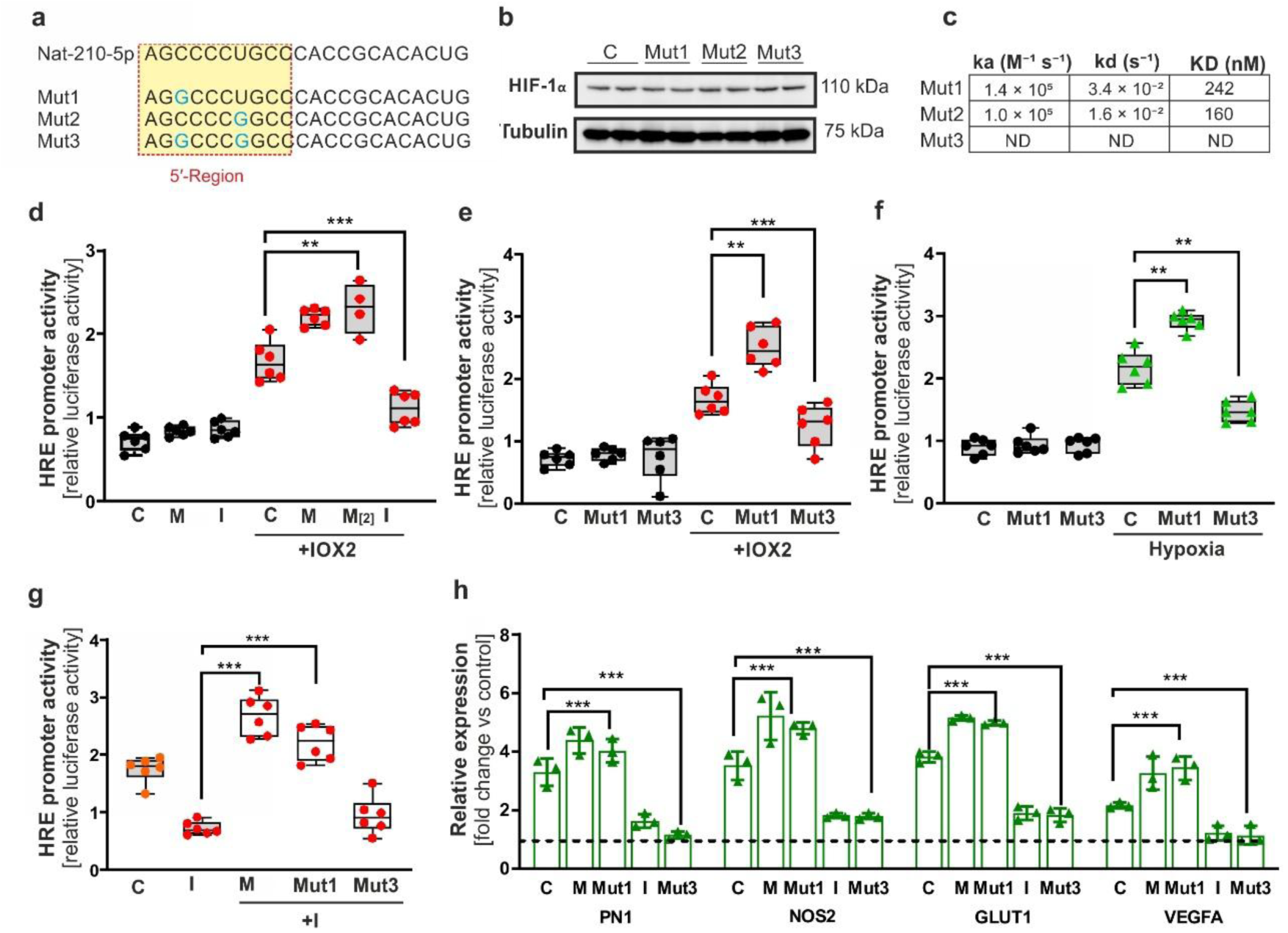
The conserved miR-210-5p 5′ motif regulates HIF-dependent transcriptional output. (a) Schematic representation of miR-210-5p and mutant variants (Mut1, Mut2, Mut3) carrying progressive nucleotide substitutions within the 5′ region. Modified nucleotides are indicated in blue. (b) Immunoblot analysis of HIF-1á protein levels following expression of miR-210-5p mimic or mutant miR-210-5p constructs in keratinocytes. Cells were transfected with control oligonucleotides (C) or mutant variants (Mut1, Mut2, Mut3; 40 nM each). Tubulin served as loading control. (c) Summary of the association rate constants (kₐ), dissociation rate constants (kd), and equilibrium dissociation constants (KD) derived from SPR analyses of miR-210-5p mutant variants interacting with recombinant HIF-1á. ND, no detectable binding. (d) HRE luciferase reporter activity following modulation of miR-210-5p levels during pharmacological HIF stabilization. Keratinocytes were co-transfected with HRE firefly luciferase and Renilla control plasmids together with control oligonucleotides (C), miR-210-5p mimics (M= 20 nM; M [2] = 40 nM), or miR-210-5p inhibitor (I = 40 nM), followed by IOX-2 treatment (50 µM, 2 h). (e) HRE luciferase reporter activity following expression of miR-210-5p mimic or mutant miR-210-5p constructs during pharmacological HIF stabilization. Keratinocytes were transfected with miR-210-5p mimic or mutant variants (Mut1 and Mut3; 40 nM each), followed by IOX-2 treatment (50 µM, 2 h). (f) HRE luciferase reporter activity following expression of miR-210-5p mimic or mutant miR-210-5p constructs under hypoxic conditions. Keratinocytes were transfected with miR-210-5p or mutant variants (Mut1 and Mut3; 40 nM each) and exposed to hypoxia (1% O₂, 24 h). (g) Sequential inhibition and reconstitution analysis of HRE luciferase activity. Endogenous miR-210-5p was inhibited using miR-210-5p inhibitor (I = 40 nM) prior to transfection with miR-210-5p mimic or mutant miR-210-5p constructs (40 nM each), followed by IOX-2 treatment (50 µM, 2 h). (h) Relative expression of PN1, NOS2, GLUT1, and VEGFA following expression of miR-210-5p mimic or mutant miR-210-5p constructs under hypoxic conditions. Keratinocytes were transfected with miR-210-5p mimic (M = 40 nM), inhibitor (I = 40 nM), or mutant variants (Mut1 and Mut3; 40 nM each) and exposed to hypoxia (1% O₂, 24 h). Gene expression was quantified by qPCR and normalized to GAPDH. Data are presented as mean ± s.e.m. Statistical significance was determined using one-way or two-way ANOVA with appropriate multiple-comparisons correction. Immunoblot and SPR data are representative of independent experiments.

To determine whether disruption of the 5′ motif affected HIF-1α stability, HIF-1α protein abundance was assessed following expression of the different miR-210-5p variants. No appreciable differences in HIF-1α protein levels were observed across constructs, indicating that the mutant series did not significantly alter HIF-1α stabilization (Fig. 3b).

Surface plasmon resonance analyses demonstrated that progressive perturbation of the conserved 5′ region markedly reduced HIF-1α binding affinity, whereas the Mut3 construct completely abolished detectable interaction (Fig. 3c). These findings identified the conserved 5′ region of miR-210-5p as a critical determinant for HIF-1α association. Because Mut2 displayed binding properties comparable to Mut1, it did not provide additional mechanistic information and was therefore not investigated further. Subsequent functional studies focused on Mut1 and the fully binding-deficient Mut3 construct.

We next investigated whether integrity of the interaction motif was required for modulation of HIF-dependent transcriptional output. HRE luciferase reporter assays demonstrated that miR-210-5p mimic (40nM) significantly enhanced HIF-dependent transcriptional output during IOX-2 treatment (Fig. 3d), whereas progressive disruption of the interaction motif reduced reporter activation in parallel with reduced HIF-1α binding (Fig. 3e). Mut1 retained partial activity, while Mut3 failed to support efficient HRE activation.

Comparable results were observed under physiological hypoxia, where disruption of the miR-210-5p 5′ motif significantly impaired HIF-dependent reporter activation (Fig. 3f). Rescue of transcriptional activity by miR-210-5p mimic but not binding-deficient variants further supported a direct functional requirement for the conserved 5′ motif (Fig. 3g). Consistent with these findings, expression of canonical HIF-responsive genes including Protease Nexin-1 (PN1), NOS2, GLUT1, and VEGFA was significantly reduced in cells expressing binding-deficient miR-210-5p variants compared with miR-210-5p mimic (Fig. 3h). Importantly, disruption of the HIF-1α interaction motif did not impair repression of the canonical cytoplasmic target ISCU (Supplementary Fig. 3b), indicating that the introduced mutations selectively disrupted HIF-1α interaction while preserving canonical miRNA activity.

Collectively, these results demonstrate that integrity of the conserved 5′ motif is required for HIF-1α interaction and potentiation of HIF-dependent transcription under hypoxic conditions.

### AGO2 forms RNA-sensitive nuclear HIF-1α complexes

Because canonical miRNA functions are frequently mediated through AGO-containing ribonucleoprotein complexes, we next investigated whether AGO2 contributes to nuclear HIF- associated assemblies. Confocal microscopy revealed increased nuclear proximity between AGO2 and HIF-1α under hypoxic conditions (Fig. 4a), with partial overlap of both signals within discrete nuclear foci (Fig. 4b).

**Figure 4.**
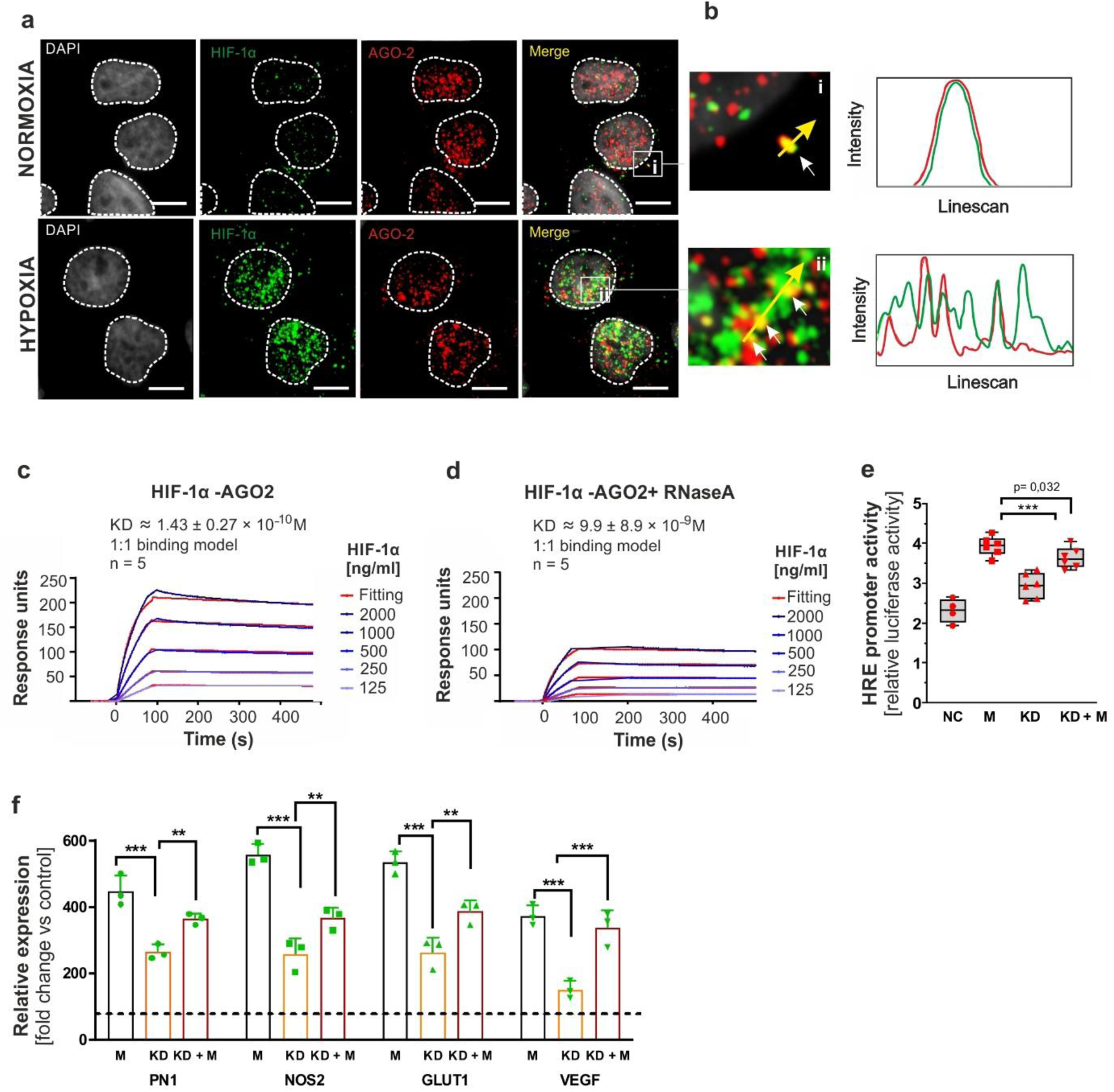
AGO2 associates with HIF-1α-containing complexes during hypoxic signaling. (a) Representative confocal microscopy images of keratinocytes cultured under normoxic conditions or exposed to hypoxia (1% O₂, 24 h). HIF-1α was detected by immunofluorescence staining (green), AGO2 by immunofluorescence staining (red), and nuclei by DAPI staining (gray). Enlarged regions corresponding to the indicated boxed areas are shown in (b). (b) Higher-magnification views and corresponding fluorescence intensity line-scan analyses of AGO2 and HIF-1á signals under normoxic (i) and hypoxic (ii) conditions. Fluorescence intensity profiles were measured along the yellow line. (c) Surface plasmon resonance (SPR) analysis of interaction between recombinant HIF-1á and AGO2. Representative sensorgrams and fitted curves derived from a 1:1 binding model are shown. Increasing concentrations of recombinant HIF-1á (125–2000 ng/ml) were injected as indicated. (d) SPR analysis of HIF-1á and AGO2 interaction following RNase A treatment. Representative sensorgrams and fitted curves derived from a 1:1 binding model are shown. Increasing concentrations of recombinant HIF-1α (125– 2000 ng/ml) were injected as indicated. (e) HRE luciferase reporter activity following AGO2 depletion and miR-210-5p overexpression during pharmacological HIF stabilization. Keratinocytes were co-transfected with HRE firefly luciferase and Renilla control plasmids together with control oligonucleotides (NC), miR-210-5p mimic (M = 40 nM), AGO2 siRNA (KD), or AGO2 siRNA together with miR-210-5p mimic (KD + M), followed by IOX-2 treatment (50 µM, 2 h). (f) Relative expression of PN1, NOS2, GLUT1, and VEGFA following AGO2 depletion and miR-210-5p overexpression under hypoxic conditions. Keratinocytes were transfected with miR-210-5p mimic (M = 40 nM), AGO2 siRNA (KD), or AGO2 siRNA together with miR-210-5p mimic (KD + M) and exposed to hypoxia (1% O₂, 24 h). Gene expression was quantified by qPCR and normalized to GAPDH. Data are presented as mean ± s.e.m. Statistical significance was determined using one-way or two-way ANOVA with appropriate multiple-comparisons correction. Immunoblot and SPR data are representative of independent experiments. Scale bars, 10 µm unless otherwise indicated.

Co-immunoprecipitation experiments demonstrated reciprocal association between endogenous AGO2 and HIF-1α in nuclear extracts obtained under hypoxic conditions (Supplementary Fig. 4a, b). Surface plasmon resonance analysis demonstrated direct interaction between recombinant HIF-1α and AGO2 (Fig. 4c), whereas RNase A treatment substantially weakened complex formation, indicating that RNA contributes to stabilization of the HIF-1α/AGO2 interaction (Fig. 4d). Despite this contribution of RNA to AGO2–HIF complex stability, direct interaction between recombinant HIF-1α and miR-210-5p was established independently by SPR and NMR analyses (Fig. 2).

To determine whether AGO2 contributes functionally to HIF transcriptional responses, HRE luciferase reporter assays were performed following AGO2 depletion and miR-210-5p overexpression. AGO2 depletion significantly reduced HIF-dependent transcriptional output, whereas co-expression of miR-210-5p partially restored reporter activation (Fig. 4e). Consistently, expression of canonical HIF target genes, including PN1, NOS2, GLUT1, and VEGFA, was reduced upon AGO2 depletion and partially rescued by miR-210-5p expression (Fig. 4f). Efficient AGO2 depletion was confirmed by immunoblot analysis (Supplementary Fig. 2b).

Collectively, these results position AGO2 as a component of nuclear HIF-associated complexes while indicating that RNA-dependent interactions contribute to hypoxic transcriptional regulation.

### miR-210-5p enhances early HIF-dependent glycolytic adaptation

Given the central role of HIF-1α in metabolic reprogramming, we next investigated whether miR-210-5p–dependent modulation of HIF activity influences cellular metabolism. Extracellular flux analysis revealed that expression of miR-210-5p mimic significantly increased extracellular acidification rates (ECAR) following IOX-2 treatment, whereas the interaction-deficient Mut3 variant failed to enhance glycolytic activity (Fig. 5a). Quantitative analysis confirmed increased basal glycolysis and glycolytic capacity in cells expressing the miR-210-5p mimic compared with Mut3-expressing cells (Fig. 5b). In contrast, oxygen consumption rates (OCR) remained largely unchanged across conditions (Supplementary Fig. 5a). Because these experiments were performed under acute HIF activation, the experimental design was intended to minimize secondary mitochondrial effects associated with prolonged canonical miR-210 activity. Accordingly, the observed metabolic changes primarily reflect enhanced early glycolytic adaptation rather than broad alterations in mitochondrial respiration.

**Figure 5.**
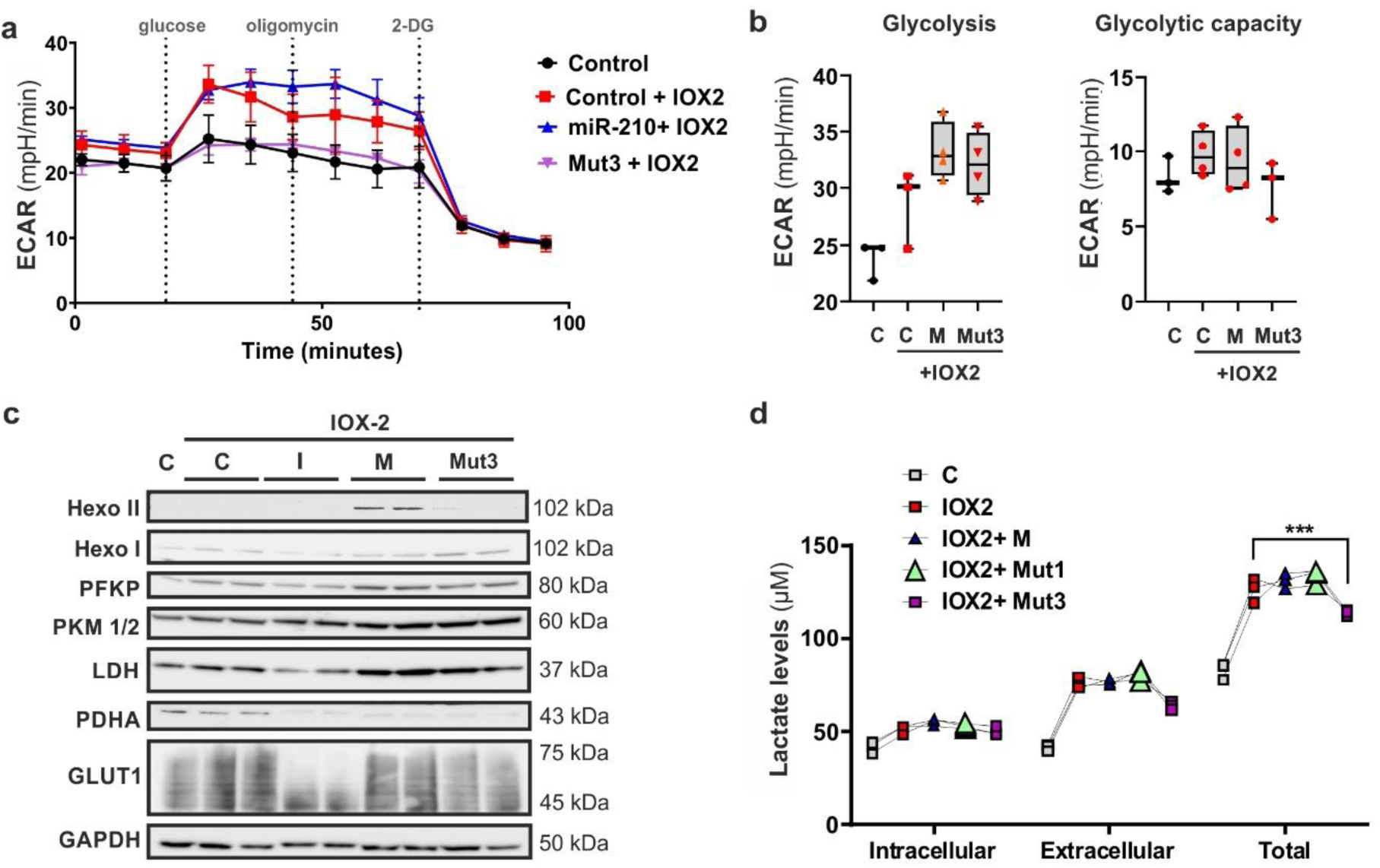
miR-210-5p enhances glycolytic adaptation under hypoxic signaling conditions. (a) Extracellular acidification rate analysis following expression of miR-210-5p mimic or mutant miR-210-5p under pharmacological HIF stabilization. Extracellular acidification rate (ECAR) traces were measured using the Seahorse XF Glycolysis Stress Test in primary human keratinocytes transfected with miR-210-5p or Mut3 constructs (40 nM). Twenty-four hours after transfection, cells were treated with IOX-2 (50 µM, 2 h) prior to and during the assay. ECAR was recorded following sequential injection of glucose, oligomycin, and 2-deoxy-D-glucose (2-DG). (b) Quantification of glycolytic parameters derived from ECAR measurements. Basal glycolysis and glycolytic capacity were calculated from ECAR traces shown in (a) using Wave software according to the manufacturer’s guidelines. (c) Immunoblot analysis of glycolysis-associated proteins following expression of miR-210-5p mimic or mutant miR-210-5p constructs. Whole-cell lysates from keratinocytes treated as in (A) were analyzed by immunoblotting for glycolysis-associated proteins including PFKP, PKM1/2, LDH, PDHA, and GLUT1. Tubulin served as loading control. (d) Quantification of intracellular and extracellular lactate levels following expression of miR-210-5p mimic or mutant miR-210-5p constructs. Total and extracellular lactate levels were measured using a luminescence-based lactate assay in keratinocytes treated as in (a). Lactate values were normalized to total protein content. Data are presented as mean ± s.e.m. from independent biological replicates. Statistical significance was determined using two-way ANOVA with appropriate multiple-comparisons correction as described in Methods.

Consistent with these metabolic changes, immunoblot analyses together with quantitative densitometric analysis demonstrated increased abundance of multiple HIF-responsive glycolytic regulators, including PFKP, PKM1/2, LDH, PDHA, and GLUT1, in cells expressing miR-210-5p but not the binding-deficient Mut3 variant (Fig. 5c, quantified on Supplementary Fig. 5b). Total and extracellular lactate production followed the same pattern and were significantly increased in miR-210-5p expressing cells (Fig. 5d).

These data indicate that miR-210-5p promotes glycolytic adaptation in proportion to its ability to interact with HIF-1α.

### Nuclear miR-210-5p colocalizes with HIF-1α in vivo

To assess the physiological relevance of the nuclear miR-210-5p/HIF-1α axis, we analyzed murine myocardium following ischemic injury (Fig. 6). Validation of the ischemia model and experimental controls are provided in Supplementary Fig. 5a-c.

**Figure 6.**
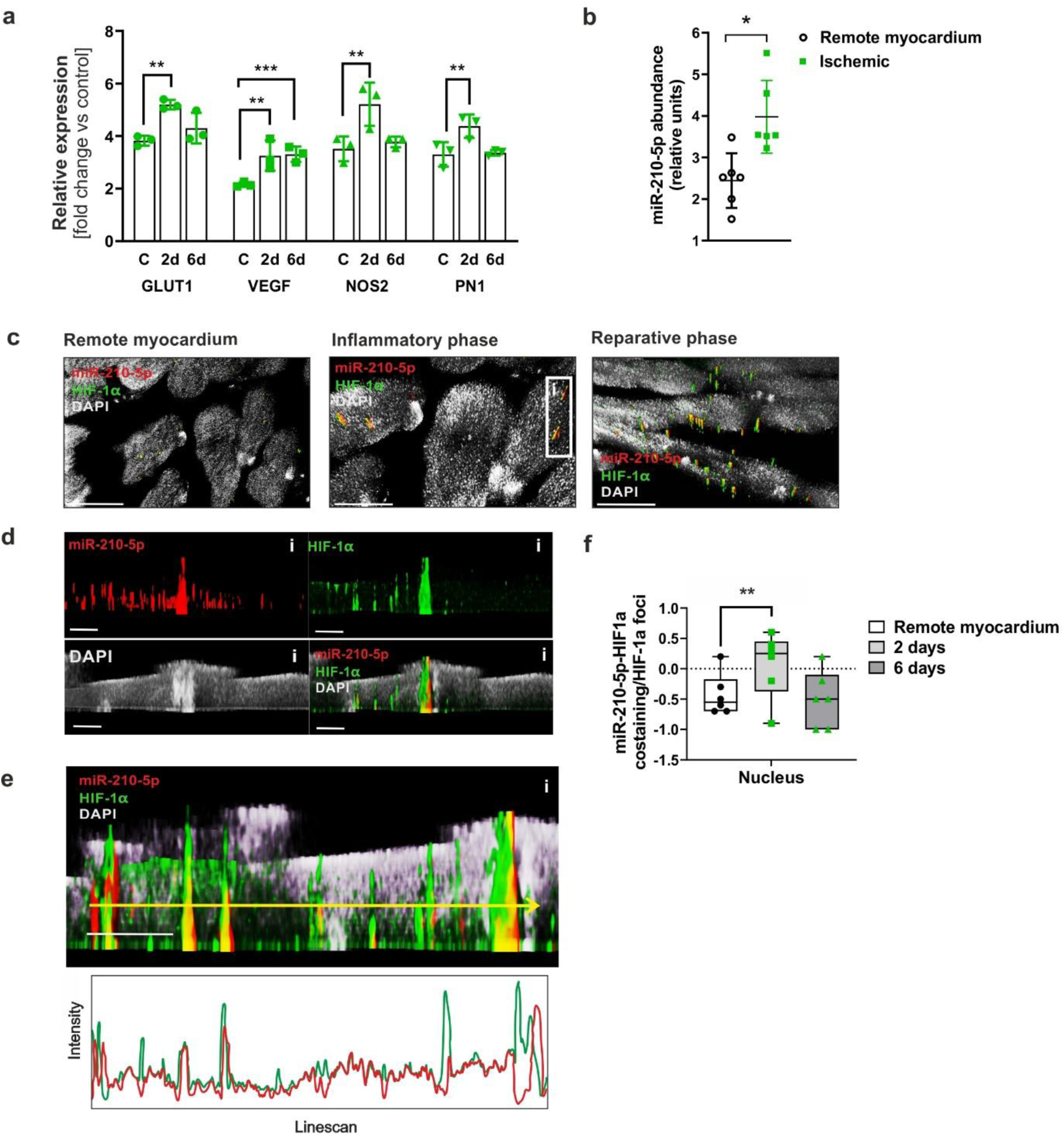
Nuclear accumulation of miR-210-5p in ischemic myocardium and spatial distribution relative to HIF-1α in vivo. (a) Relative expression of GLUT1, VEGFA, NOS2, and PN1 in myocardial tissue collected 2 and 6 days following ischemic injury. Gene expression was quantified by qPCR and normalized to GAPDH. Expression levels are shown relative to remote myocardium. (b) Relative miR-210-5p abundance in remote and ischemic myocardium following myocardial infarction. miR-210-5p levels were quantified by qPCR and normalized to sno202. (c) Representative RNA-FISH and immunofluorescence images of myocardial tissue sections obtained from remote myocardium, ischemic myocardium at day 2, and ischemic myocardium at day 6 following injury. miR-210-5p was detected by RNA-FISH (red), HIF-1α by immunofluorescence staining (green), and nuclei by DAPI staining (gray). Enlarged regions are shown for the indicated boxed area. (d) Representative orthogonal fluorescence views of miR-210-5p, HIF-1α, and DAPI signals within myocardial tissue sections collected at day 2 following ischemic injury. Individual fluorescence channels and merged images are shown. (e) Representative three-dimensional reconstruction and fluorescence intensity line-scan analysis of miR-210-5p and HIF-1á signals within myocardial tissue sections collected at day 2 following ischemic injury. Fluorescence intensity profiles were measured along the yellow line. (f) Quantification of Pearson’s correlation coefficients between miR-210-5p and HIF-1α fluorescence signals within nuclear regions from remote and ischemic myocardium collected at day 2 and day 6 following injury. Data are presented as mean ± s.e.m. Statistical significance was determined using unpaired two-tailed Student’s t-test or one-way ANOVA with appropriate multiple-comparisons correction. Scale bars, 10 µm unless otherwise indicated.

Expression of canonical HIF-responsive genes, including GLUT1, VEGFA, NOS2, and PN1, was increased during post-ischemic remodeling phases (Fig. 6a). Consistently, miR-210-5p abundance was significantly elevated in ischemic compared with remote myocardium (Fig. 6b). RNA-FISH analyses demonstrated increased miR-210-5p expression within ischemic compared with remote myocardium (Fig. 6c). High-resolution confocal microscopy further revealed prominent nuclear miR-210-5p puncta in close spatial proximity to HIF-1α within ischemic myocardium (Fig. 6d). Orthogonal views and three-dimensional reconstructions confirmed focal nuclear overlap between miR-210-5p and HIF-1α signals within cardiomyocyte nuclei (Fig. 6e).

Quantitative colocalization analyses demonstrated significantly increased nuclear miR-210-5p/HIF-1α spatial association following ischemic injury compared with remote myocardium (Fig. 6f). Consistent with these findings, re-analysis of publicly available single-nucleus RNA-sequencing datasets revealed transient activation of canonical HIF-responsive transcriptional programs during early post-ischemic phases (Supplementary Fig. 6d).

Together, these findings support the physiological relevance of the nuclear miR-210-5p/HIF-1α regulatory axis in ischemic tissue, and indicate that hypoxia-associated nuclear accumulation of miR-210-5p occurs in vivo during pathological oxygen deprivation. These observations prompted us to consider how direct interaction between miR-210-5p and HIF-1α may contribute to transcriptional regulation during hypoxic adaptation.

## Discussion

The mechanisms that determine the magnitude and specificity of HIF-dependent transcriptional responses remain incompletely understood. Here, we identify a previously unrecognized nuclear function of the hypoxamiR miR-210-5p by demonstrating that it directly associates with HIF-1α through a conserved 5′ motif-dependent RNA–protein interaction and enhances HIF-dependent transcription independently of its established post-transcriptional activities (Fig. 7). These findings reveal an additional layer of hypoxic gene regulation and support a model in which mature nuclear miRNAs can directly modulate transcription factor function during cellular adaptation to hypoxic stress.

**Figure 7.**
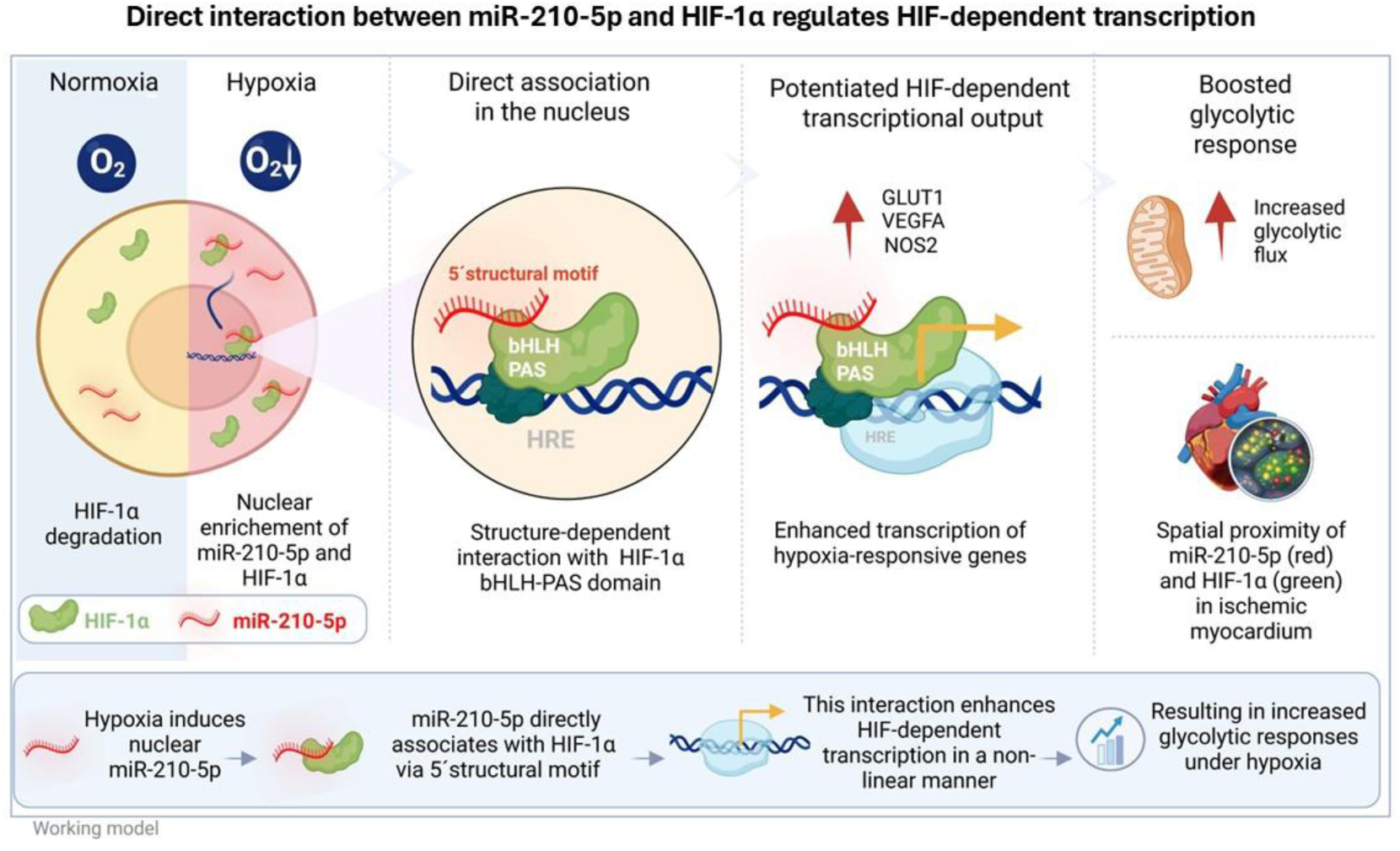
Proposed working model of the miR-210-5p–HIF-1α regulatory axis under hypoxia. Schematic summary of the proposed mechanism derived from the findings presented in this study. Under normoxic conditions, HIF-1α undergoes rapid degradation, whereas hypoxia promotes HIF-1α stabilization and nuclear enrichment of mature miR-210-5p. Within the nucleus, miR-210-5p associates with the HIF-1α bHLH–PAS domain through a conserved 5′ interaction motif. This interaction is associated with enhanced HIF-dependent transcriptional output and increased expression of hypoxia-responsive genes, including glycolysis-associated targets. The relationship between miR-210-5p–HIF-1α interaction and transcriptional output is consistent with a non-linear mode of transcriptional potentiation. Increased glycolytic responses and spatial proximity between nuclear miR-210-5p and HIF-1α in ischemic myocardium support the physiological relevance of this proposed regulatory axis. Created with BioRender.com

Hypoxic signaling induced rapid and HIF-dependent nuclear enrichment of mature miR-210-5p across multiple cellular contexts.^19^ This redistribution occurred within a timeframe compatible with early transcriptional responses and was reversible following withdrawal of HIF-stabilizing stimuli, indicating that nuclear localization represents a dynamic component of the hypoxic response. Unlike canonical mechanisms regulating HIF activity, miR-210-5p associated with the HIF-1α bHLH–PAS domain without measurably altering HIF-1α abundance or stability. These observations suggest that nuclear miR-210-5p modulates the transcriptional competence of stabilized HIF-1α rather than its accumulation and propose subcellular redistribution of mature miRNAs as an additional regulatory layer within hypoxic signaling.

A central finding of this study is that a mature hypoxia-induced miRNA can directly engage a transcription factor through a conserved structural element. Although nuclear localization has been described for several miRNAs, the molecular basis by which they selectively regulate transcriptional machinery remains poorly understood.^12,14,20–22^ Our data demonstrate that a conserved 5′ motif within miR-210-5p is required for HIF-1α interaction and transcriptional potentiation while remaining dispensable for repression of the canonical cytoplasmic target ISCU. Together with the concordance between predicted structural perturbation, reduced HIF-1α interaction in biophysical assays, and diminished transcriptional activity, these findings demonstrate that the nuclear and cytoplasmic activities of miR-210-5p can be experimentally separated and establish direct RNA–protein interaction as a distinct mechanism of nuclear miRNA function.^23^

These observations complement, rather than replace, the established metabolic functions of miR-210. Previous studies demonstrated that miR-210 regulates mitochondrial metabolism through repression of targets including ISCU and SDHD, thereby influencing reactive oxygen species production and HIF signaling during prolonged hypoxic adaptation.^9,10^ Such mechanisms are likely to contribute to the broader physiological functions of miR-210 but cannot fully explain the rapid transcriptional effects observed here. Nuclear accumulation occurred shortly after HIF activation, transcriptional potentiation was observed without substantial changes in HIF-1α abundance, and selective disruption of the HIF-interacting structural motif abolished transcriptional enhancement while preserving repression of ISCU. Furthermore, OCR measurements performed under acute HIF activation conditions revealed minimal changes in mitochondrial respiration, consistent with the experimental design, which was intended to minimize secondary metabolic consequences of prolonged repression of canonical miR-210 targets. Together, these findings support the existence of a mechanistically distinct nuclear activity operating alongside the established cytoplasmic functions of miR-210 and suggest that this regulatory axis may be particularly important during the early phases of hypoxic adaptation.

Previous studies have implicated nuclear non-coding RNAs, including miRNAs and lncRNAs, in transcriptional regulation through promoter-associated mechanisms, chromatin-associated regulatory complexes, and RNA–DNA triplex formation.^24–26^ Our findings instead support a distinct mechanism in which a mature miRNA directly engages a transcription factor through a motif-dependent RNA–protein interaction. Nuclear AGO-containing complexes have been implicated in transcriptional regulation through interactions with RNA polymerase II, chromatin-associated proteins, and epigenetic regulators.^20,27^ Consistent with this framework, AGO2 contributed to HIF-dependent transcriptional responses and was detected within nuclear HIF-1α-containing complexes. However, direct interaction between miR-210-5p and HIF-1α was readily detected in purified biophysical assays performed in the absence of AGO2, demonstrating that AGO2 is not required for physical association between the two molecules. These findings support a model in which direct miRNA–transcription factor interaction and AGO2-dependent regulatory pathways coexist as partially separable mechanisms controlling hypoxic gene expression.^28–31^ Whether AGO2 primarily stabilizes these complexes or facilitates recruitment of additional transcriptional cofactors remains an important question for future investigation.^28^

Functionally, miR-210-5p enhanced glycolytic adaptation and increased expression of selected HIF-responsive genes without broadly altering mitochondrial respiration. These findings are consistent with modulation of HIF-dependent transcriptional output rather than binary activation of hypoxic signaling and may help explain how cells generate graded transcriptional responses despite comparable levels of stabilized HIF-1α. Notably, the relationship between RNA–protein interaction and transcriptional activity appeared non-linear, as variants retaining partial HIF-1α binding also preserved intermediate transcriptional activity. These observations suggest that relatively modest differences in RNA–protein interaction strength may be sufficient to fine-tune transcriptional responses under hypoxic conditions.

A limited analysis of miR-21-5p demonstrated HIF-dependent nuclear enrichment together with direct interaction with HIF-1α and enhanced HIF-dependent transcription (Supplementary Fig. 7). These observations suggest that direct transcription factor engagement may not be unique to miR-210-5p. However, in contrast to miR-210-5p, the molecular determinants underlying the miR-21-5p interaction were not investigated in detail, and no conclusions can therefore be drawn regarding the underlying mechanism. Future studies examining additional hypoxia-associated miRNAs will be required to determine whether direct RNA–transcription factor interactions represent a broader principle of nuclear hypoxamiR biology.

The physiological relevance of this regulatory axis is supported by the accumulation of nuclear miR-210-5p and its close spatial proximity to HIF-1α within ischemic myocardium.^32,33^ Although these observations do not establish a causal interaction in vivo, they demonstrate that the nuclear redistribution observed in cultured cells is maintained under pathological hypoxic conditions. The concordance between cellular, biophysical, functional, and tissue-level observations therefore supports the physiological relevance of the miR-210-5p–HIF-1α interaction.

Several mechanistic questions remain unresolved. Although our data establish the specificity and functional relevance of the miR-210-5p–HIF-1α interaction, the downstream molecular consequences of this association remain to be defined. In particular, whether miR-210-5p influences HIF chromatin occupancy, transcriptional complex stability, cofactor recruitment, or promoter residence time will require further investigation.^12^ Likewise, high-resolution structural characterization of the interaction interface and direct in vivo disruption of the RNA–protein interaction motif represent important future objectives. The present study was designed to establish the existence and functional significance of this regulatory axis rather than to fully resolve its downstream chromatin-associated mechanisms.

In summary, our findings identify miR-210-5p as a nuclear regulator of HIF-dependent transcription that acts through direct association with HIF-1α. Beyond its established role in post-transcriptional gene regulation, miR-210-5p modulates HIF-dependent transcriptional output during hypoxic adaptation through a conserved 5′ motif-dependent RNA–protein interaction (Fig. 7). Together with our observations on miR-21-5p, these findings raise the possibility that direct engagement of transcription factors represents a broader property of selected nuclear hypoxamiRs. More broadly, our findings extend current models of nuclear miRNA biology by identifying direct interactions with transcriptional regulatory proteins as an additional mechanism through which mature miRNAs can regulate adaptive gene expression. This work establishes a conceptual framework for investigating nuclear miRNA–transcription factor interactions as a previously unrecognized layer of transcriptional regulation and provides a foundation for exploring how these mechanisms may be harnessed to modulate hypoxia-driven gene expression in human disease.

## Material and Methods

### Cell culture and hypoxic stimulation

HEK293 cells, H9C2 cardiomyoblasts, and 4T1 breast cancer cells were maintained in DMEM, whereas normal human epidermal keratinocytes (NHEK) were cultured in keratinocyte growth medium (Lonza). Human umbilical vein endothelial cells (HUVECs) were grown in endothelial growth medium (Lonza). All media (excluding NHEK) were supplemented with 10% fetal bovine serum, 1% penicillin–streptomycin, and 2 mM L-glutamine unless otherwise specified by the manufacturer. Cells were incubated at 37 °C in a humidified atmosphere containing 5% CO2.

Hypoxic stimulation was performed at 1% O2, 5% CO2, and 94% N2 using a ProOx C21 hypoxia chamber (BioSpherix). Normoxic control cells were maintained at atmospheric oxygen levels (21% O2). Where indicated, chemical stabilization of HIF was achieved by treatment with IOX-2 (50 µM) for the indicated time points. In selected experiments, treatment with ivermectin, an inhibitor of importin α/β-mediated nuclear import (IVE) was used (5 µM).

For targeted depletion of HIF-1α, RNA interference was carried out using SMARTpool siRNA directed against HIF-1α (Dharmacon, Lafayette, CO, USA), while non-targeting control siRNA served as negative control. Reverse transfection was performed using Lipofectamine RNAiMAX (Thermo Fisher Scientific) according to the manufacturer’s instructions.

For miRNA functional analyses, miRCURY LNA™ miRNA mimics and inhibitors targeting miR-210-5p (Qiagen, Hilden, Germany), together with corresponding negative control oligonucleotides, were used. miRNA oligonucleotides were transfected at a final concentration of 20 nM using Lipofectamine 2000 (Invitrogen, Waltham, MA, USA) following established protocols. A complete list of all siRNAs and miRNA reagents, including sequences and catalogue numbers, is provided in Table S1.

### miRNA mimics, inhibitors, and mutant design

Chemically synthesized miR-210-5p mimics and locked nucleic acid (LNA)-based inhibitors (Qiagen) were transfected using Lipofectamine 2000 (Thermo Fisher Scientific) at final concentrations of 20–40 nM, as indicated for individual experiments. Mutant variants of miR-210-5p (Mut1–Mut3) carrying progressive nucleotide substitutions within the conserved 5′ region were custom synthesized by Eurofins Genomics (Ebersberg, Germany). Mutations were designed to progressively disrupt the predicted native 5′ interaction motif while preserving overall miRNA length and general sequence composition. Mutant oligonucleotides were transfected under the same experimental conditions as miR-210-5p mimic.

RNA secondary structure predictions for native and mutant miRNAs were generated using RNAfold (ViennaRNA Package).

### RNA Isolation and Quantitative PCR (qPCR) Analysis

Total RNA was isolated from cells using the RNeasy Mini Kit (Qiagen, Hilden, Germany) according to the manufacturer’s instructions. To eliminate genomic DNA contamination, on-column DNase digestion was performed using the RNase-free DNase Set (Qiagen). For mRNA analysis, cDNA synthesis was carried out using the QuantiTect Reverse Transcription Kit (Qiagen). Quantitative PCR was performed on a QuantStudio 5 Real-Time PCR System (Thermo Fisher Scientific) using the QuantiTect SYBR Green PCR Kit (Qiagen) or GoTaq® qPCR Master Mix (Promega), as indicated. For miRNA expression analysis from total RNA, reverse transcription and qPCR were performed using either the miRCURY LNA PCR System (Qiagen) or miScript technology (Qiagen), according to the manufacturer’s protocols.

For nuclear and cytoplasmic RNA analyses, subcellular RNA fractions were obtained using the Cytoplasmic and Nuclear RNA Purification Kit (Norgen Biotek). miRNA expression from fractionated RNA samples was quantified using miScript technology following the manufacturer’s instructions. Fraction purity was verified for each experiment.

Relative miRNA expression levels were normalized to stable small RNA reference genes (sno202, snord61, snord72, U2, or U6b), as indicated for each experiment. mRNA expression levels were normalized to GAPDH. Relative expression values were calculated using the ΔCt or ΔΔCt method and scaled to the sample with the lowest expression level. Primer sequences and catalogue numbers are provided in Table S3.

### Protein Isolation and Analysis

Approximately 1.5 × 10⁶ cells per sample were lysed in RIPA buffer containing 50 mM Tris (pH 8.0; Merck, Darmstadt, Germany), 150 mM NaCl (Merck), 0.1% (w/v) SDS (Carl Roth, Karlsruhe, Germany), 5 mM EDTA (Merck), and 0.5% (w/v) sodium deoxycholate (Sigma-Aldrich), freshly supplemented with a protease inhibitor cocktail (Sigma-Aldrich). Cell disruption was performed using a tissue lyser, followed by two centrifugation steps to remove lipids and cellular debris. Protein concentrations were determined using the Pierce BCA Assay Kit (Thermo Fisher Scientific).

For immunoblotting, 15–30 µg of protein per sample was denatured in the presence of DTT (Sigma-Aldrich) at 95 °C for 3 min and separated on NuPAGE 4–12% Bis-Tris gels (Thermo Fisher Scientific). Proteins were transferred onto 0.2 µm PVDF membranes (Bio-Rad, Hercules, CA, USA) using the Trans-Blot Turbo™ transfer system (Bio-Rad) at 25 V and 1.3 A for 7 min. Membranes were blocked for 1 h at room temperature in 5% skim milk prepared in TBS-T.

Primary antibody incubations were carried out overnight at 4 °C using antibodies diluted 1:1000 in 5% skim milk (Table S3). Membranes were subsequently washed with TBS-T (Tris-buffered saline containing 0.1% Tween 20) and incubated with HRP-conjugated secondary antibodies (Cell Signaling Technology, Danvers, MA, USA) at a dilution of 1: 10,000 for 1 h at room temperature. Protein signals were detected using SuperSignal™ West Pico PLUS Chemiluminescent Substrate (Thermo Fisher Scientific) and visualized with a ChemiDoc MP Imaging System (Bio-Rad). All original, uncropped immunoblots are provided in the accompanying Source Data file.

### Recombinant HIF-1α constructs

Recombinant HIF-1α fragments were designed based on annotated domain organization together with intrinsic disorder prediction to generate soluble protein constructs suitable for biophysical studies. Constructs comprising amino acid residues 1–238, 1–358, 238–358 and 786–826 of human HIF-1α were cloned, recombinantly expressed in *Escherichia coli* and purified by affinity chromatography followed by size-exclusion chromatography. For NMR spectroscopy, uniformly ^15N-labelled HIF-1α (1–238) was produced in M9 minimal medium supplemented with ^15NH4Cl as the sole nitrogen source. Purified proteins were concentrated and exchanged into NMR buffer before measurements.

### Surface Plasmon Resonance (SPR)

Experiments were performed using a Biacore X100 instrument, SA and C1 chips (Cytiva, Freiburg, Germany). Biotinylated native and mutated miR-210-5p, and miR-21-5p were immobilized on a SA chip at the density of approximately 300 Resonance Units (RU). AGO2 was immobilized by thiol coupling on a C1 chip at a density of 740 RU. Recombinant HIF-1a and truncation constructs were injected in HBS-EP⁺ buffer at a flow rate of 60 µL/min and 90 µL/min, respectively. In the case of the RNAse treatment assay, RNAse A (Cell Signaling Technology, Frankfurt am Main, Germany) was added in the HBS-EP+ buffer at the final concentration of 1µg/mL. Association and dissociation phases were recorded for 90 s and 300 s, except for truncated constructs that were perfused during 60 s. Sensor surfaces were regenerated with two 60 s pulses of 30 mM NaOH and 2 M NaCl. Binding data were fitted using a 1:1 Langmuir interaction model in the Biacore X100 Evaluation Software 2.0.1 Plus (Cytiva).

### Nuclear magnetic resonance (NMR) spectroscopy

NMR spectroscopy was performed to independently validate the direct interaction between HIF-1α and miR-210-5p. Uniformly ^15N-labelled recombinant HIF-1α(1–238), comprising the structured N-terminal bHLH–PAS region identified by domain architecture and intrinsic disorder prediction, was used for all NMR measurements.

Protein samples were prepared at approximately 100–200 μM in NMR buffer containing 20 mM sodium phosphate (pH 6.8), 150 mM NaCl, 1 mM DTT and 10% D2O. Synthetic mature miR-210-5p was added immediately before data acquisition.

Two-dimensional ^1H–^15N HSQC spectra were recorded on a Bruker Avance spectrometer equipped with a cryogenic probe. Spectra of free HIF-1α(1–238) were compared with spectra recorded after addition of miR-210-5p to assess RNA-induced spectral changes. In addition, one-dimensional ^1H NMR spectra of the imino proton region were acquired to monitor the RNA signal in the presence of HIF-1α.

NMR data were processed using NMRPipe and analysed with NMRView.

### Seahorse extracellular flux analysis (glycolysis stress-test)

Glycolytic function and mitochondrial respiration were assessed using the Seahorse XF Glycolysis Stress Test (Agilent Technologies), with simultaneous measurement of extracellular acidification rate (ECAR) and oxygen consumption rate (OCR). Keratinocytes were seeded at a density of 10,000 cells per well in XF cell culture microplates and allowed to attach and equilibrate for 48 h prior to the assay. On the day of measurement, cells were washed and incubated in Seahorse XF Base Medium (Agilent) supplemented with 2 mM L-glutamine, adjusted to pH 7.4 at 37°C, and lacking glucose and bicarbonate. Plates were equilibrated for 45–60 min in a non-CO₂ incubator at 37°C prior to analysis.

Where indicated, IOX-2 was added directly to the assay medium and maintained throughout the measurement to induce chemical hypoxia. Baseline ECAR and OCR were recorded under glucose-free conditions. Glycolysis was initiated by injection of glucose (10 mM), followed by oligomycin (1 µM) to inhibit mitochondrial ATP synthase and reveal maximal glycolytic capacity. Glycolysis was subsequently inhibited by injection of 2-deoxy-D-glucose (2-DG; 50 mM) to confirm glycolytic-dependent ECAR.

ECAR and OCR measurements were recorded using standard mixing and measurement cycles according to the manufacturer’s instructions. Data were normalized to cell number and analyzed using Wave software (Agilent). Glycolytic parameters were calculated as defined by the Glycolysis Stress Test assay guidelines.

### Lactate measurements

Lactate levels were quantified using the Lactate-Glo™ Assay (Promega) according to the manufacturer’s instructions. For extracellular lactate measurements, culture supernatants were collected at the indicated time points and cleared by centrifugation at 1,000 × g for 5 min to remove cellular debris. Samples were diluted in assay buffer where necessary to ensure measurements within the linear range of the assay. For intracellular lactate measurements, cells were rapidly washed twice with ice-cold PBS to remove residual extracellular lactate and immediately lysed in lactate assay lysis buffer on ice. Lysates were clarified by centrifugation at 12,000 × g for 10 min at 4°C, and supernatants were used for analysis. Total lactate was determined by combining extracellular lactate from culture supernatants with intracellular lactate from matched cell lysates derived from the same experimental condition and normalized sample input. For all measurements, an equal volume of Lactate Detection Reagent was added to samples in white 96-well plates, followed by incubation for 60 min at room temperature. Luminescence was recorded using a microplate luminometer. Lactate concentrations were calculated from a standard curve generated using freshly prepared lactate standards. Extracellular lactate was normalized to cell number or total protein content, intracellular lactate was normalized to total protein content, and total lactate was expressed as the sum of normalized extracellular and intracellular values.

### HRE promoter reporter assay

Cells were co-transfected with an HRE-driven Firefly luciferase reporter plasmid and a constitutively expressed Renilla luciferase plasmid (internal control). After 24 h, cells were exposed to hypoxia (1% O**₂**, 24 h) or treated with the PHD inhibitor IOX-2 (50 µM, 2 h). Cells were lysed and Firefly and Renilla activities were quantified using the Dual-Luciferase Reporter Assay System (Promega) according to the manufacturer’s instructions. Firefly luciferase activity was normalized to Renilla luciferase activity, and data are presented as fold change relative to normoxic/vehicle controls.

### Immunofluorescence analysis

The cells were seeded in glass bottom chamber 8-well slides (Ibidi) 24 h before experimental treatments. Upon completion of treatments, cells were fixed with 4% paraformaldehyde (PFA) for 30 min and permeabilized with 0.1% Triton X-100 in PBS/BSA 1% for 30 min at room temperature. The slides were subsequently incubated with primary antibodies (Table S2) overnight at 4°C. Nonspecific isotype antibodies were used as negative controls. The slides were embedded in Prolong® Diamond antifade mountant (ThermoFisher Scientific) with added 4’,6-diamidino-2-phenylindole (DAPI) to counterstain nuclei.

Digital images were acquired using three-dimensional confocal laser scanning microscopy (CLSM) was performed with a Leica SP8 3X microscope equipped with a 100xNA1.40 (Leica) oil immersion objective. Optical zoom was used where applicable.

Image reconstructions were performed using the LAS X software package v.3.0.2 (Leica) and deconvolution was applied in combination with the Huygens Professional software package v.19.10 (Scientific Volume) using the unsupervised CMLE (for CLSM) or wide field (for standard immunofluorescence) algorithms. Multiple fields were acquired to image an average number of 100 cells/well, and multiple wells per experimental condition were performed in independent experiments.

### RNA-FISH and in situ PCR for miR-210-5p

For *in vitro* studies on keratinocytes, fluorescence *in situ* hybridization (FISH) and sequential branched-DNA (bDNA) amplification for single-RNA molecule sensitivity along with protein immunostaining were performed employing a viewRNA cell plus kit (Life Technology) with some modifications to the supplier’s protocol. Briefly, cells were fixed with Cell Plus fixation solution containing RNase inhibitor for 1 h, then incubated with a miR-210-5p probe (1:50 for 2 h at 40±1°C). The probe was amplified with 2x2-h incubation steps with specific Amplifier Mixes and subsequently labeled with an Alexa546 fluorochrome. Cells were then fixed again for 30 min, blocked with Cell Plus Blocking solution containing RNase inhibitors for 30 min and incubated overnight at 4°C with an antibody against HIF-1α (Cell Signaling Technology). Images were acquired by CLSM as described above. FISH was also performed on OCT-embedded murine specimens following the same protocol after having performed antigen retrieval using standard protocols. Colocalization analyses were performed on z-stacks using identical acquisition and thresholding parameters across conditions.

### Mouse model of myocardial infarction

As mice do not develop reproducible spontaneous myocardial infarction, vascular occlusion was mechanically induced by ligation of the left anterior descending coronary artery (LAD).

Surgery was performed under triple-combination anesthesia consisting of midazolam (5 mg/kg), medetomidine (0.5 mg/kg), and fentanyl (0.05 mg/kg). Mice were placed in the supine position, and the trachea was intubated for mechanical ventilation (120 strokes/min, 200 µl tidal volume; MiniVent type 845, Hugo Sachs Elektronik–Harvard Apparatus). Animals were then carefully positioned in right lateral decubitus, and a left-sided thoracotomy was performed between the third and fourth intercostal spaces. Tissue and muscle were dissected carefully using cautery to minimize bleeding. The thoracic cavity was opened, and the heart was exposed without contacting the lungs with sharp instruments.

The left anterior descending coronary artery (LAD) was permanently ligated at a proximal position using a single 7-0 silk suture (Covidien, cat. no. VS-809). Successful occlusion was confirmed by immediate blanching of the left ventricular myocardium and later verified by visual inspection during postmortem dissection. The chest wall was closed using the same 7-0 silk suture, and the skin was closed with a 5-0 silk suture (Seraflex, cat. no. DSS-13). Mice were allowed to recover gradually while remaining on mechanical ventilation.

Following surgery, anesthesia was reversed by intraperitoneal administration of flumazenil (0.5 mg/kg), atipamezole (2.5 mg/kg), and naloxone (1.2 mg/kg). Postoperative analgesia was provided by subcutaneous injection of buprenorphine (0.05 mg/kg) every 4-6 h initially, depending on pain severity and expected duration.

At the indicated time points, hearts were harvested and fixed overnight in 4% paraformaldehyde, transferred to 30% sucrose for 24 h, embedded in OCT, and sectioned at 4 µm thickness. All animal experiments were approved by the relevant institutional authorities and conducted in accordance with national guidelines.

### Immunoprecipitation (IP) and RNA-IP

Protein G–conjugated magnetic beads (Dynabeads® Protein G, Invitrogen) were equilibrated and incubated with anti–HIF-1α antibody (4 µg) or matched isotype control antibodies (Table S2) for 30 min at room temperature. Antibody-loaded beads were subsequently combined with nuclear or cytoplasmic keratinocyte extracts and rotated overnight (10–12 h) at 4 °C. Following extensive washing (five times), immune complexes were resuspended in RIP buffer containing 50 mM Tris-HCl, 150 mM NaCl, 10 mM MgCl₂, 0.005% NP-40, 5 mM DTT, 5 mM EDTA, and RNase inhibitor (250 U/mL). To assess non-associated RNA, an aliquot of the initial wash fraction was collected and processed separately for RNA isolation. RNA bound to immunoprecipitated complexes was purified using the miRNeasy Mini Kit (Qiagen) according to standard protocols. Enrichment of specific miRNAs was determined by quantitative PCR employing miScript assays (Qiagen), normalized to corresponding input samples, and corrected for background signals obtained from IgG controls.

### Statistical analysis

Unless otherwise indicated, all experiments were performed using at least three independent biological replicates unless otherwise indicated. Data are presented as mean ± standard error of the mean (s.e.m.). Statistical analyses were performed using GraphPad Prism (GraphPad Software).

Comparisons between two groups were analyzed using unpaired two-tailed Student’s *t*-tests. Comparisons involving multiple groups were analyzed using one-way or two-way analysis of variance (ANOVA) followed by Tukey’s or Sidak’s multiple-comparison test, as indicated in the corresponding figure legends.

For fluorescence colocalization analyses, Pearson’s correlation coefficients were calculated from individual nuclei using ImageJ/Fiji. For luciferase reporter assays, firefly luciferase activity was normalized to Renilla luciferase activity. *P* < 0.05 was considered statistically significant.

## Acknowledgments

The authors would like to thank all members of the Bartelt Lab for their support and insightful feedback throughout this project. We also gratefully acknowledge Thomas Pitsch for his excellent technical assistance.

## Author contributions

X.B. Designed, performed experiments, and analyzed data.

A.M., B.F., L.M., H.J., P.H., and J.C. performed experiments and analyzed data.

M.S., A.B., C.R., and C.W. analyzed data, provided supervision, and intellectual input.

V.E. Conceived, designed, performed experiments, analyzed data, supervised the study, and wrote the manuscript.

## Data and materials availability

All data associated with this study are available in the main text, the Supplementary Materials, and the accompanying Source Data file containing the full, uncropped immunoblot images.

