## Supplementary Information for "Direct interaction between miR-210-5p and HIF-1α regulates HIF-dependent transcription"

a

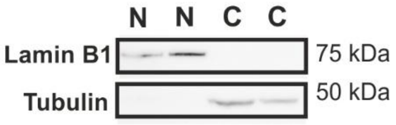


**Supplementary Figure 1. Validation of nuclear fractionation, HIF-1α protein levels, nuclear miR-210 induction kinetics, and preservation of canonical ISCU repression.**

**(a)** Immunoblot analysis of nuclear (N) and cytosolic (C) fractions from primary human keratinocytes following IOX-2 treatment (50 μM, 2 h) or under control conditions. Fraction purity was assessed using Lamin B1 as a nuclear marker and Tubulin as a cytosolic marker.

**(b)** qPCR analysis of ISCU mRNA levels in keratinocytes expressing native or mutant miR-210-5p constructs (Mut1 and Mut3; 40 nM, 24 h). Mut1 and Mut3 were selected as representative partial-binding and binding-deficient variants. Expression was normalized to GAPDH and shown relative to control.

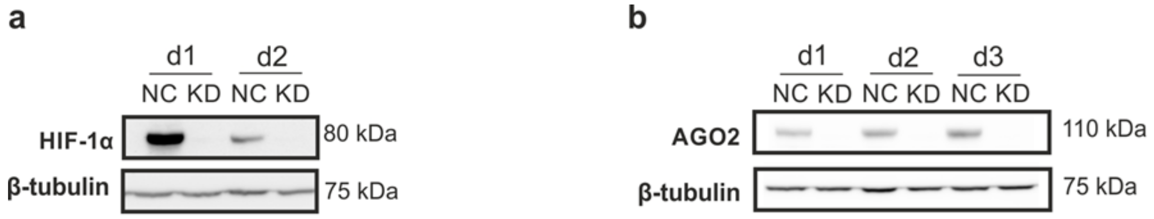


**Supplementary Figure 2. Validation of HIF-1α and AGO2 depletion.**

(a) Validation of HIF-1α depletion. Keratinocytes were transfected with control (NC) or HIF-1α-targeting siRNA (KD) and harvested at the indicated time points (d1–d2). HIF-1α protein levels were assessed by immunoblotting. Tubulin served as a loading control.

(b) Validation of AGO2 depletion. Keratinocytes were transfected with control (NC) or AGO2-targeting siRNA (KD) and harvested at the indicated time points (d1–d3). AGO2 protein levels were assessed by immunoblotting. Tubulin served as a loading control.

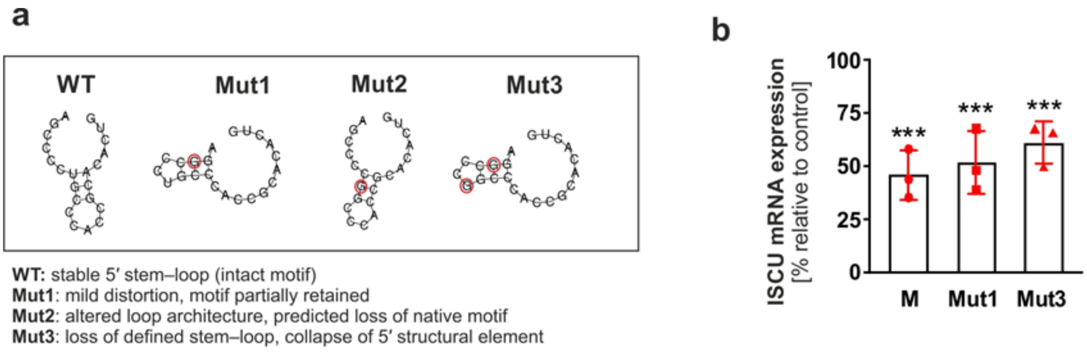


**Supplementary Figure 3. Predicted secondary structures of native and mutant miR-210-5p variants and preservation of canonical ISCU repression.**

(a) Predicted secondary structures of native miR-210-5p and mutant variants (Mut1, Mut2, and Mut3) generated using RNAfold (ViennaRNA Package). Structures represent minimum free-energy predictions based on the corresponding RNA sequences. Nucleotide substitutions introduced into the mutant variants are indicated in Fig. 3a**.**

(b) qPCR analysis of ISCU mRNA expression in keratinocytes transfected with native miR-210-5p or mutant constructs (Mut1 and Mut3; 40 nM, 24 h). Mut1 and Mut3 were selected as representative partial-binding and binding-deficient variants, respectively. ISCU expression was normalized to GAPDH and is presented relative to control cells.

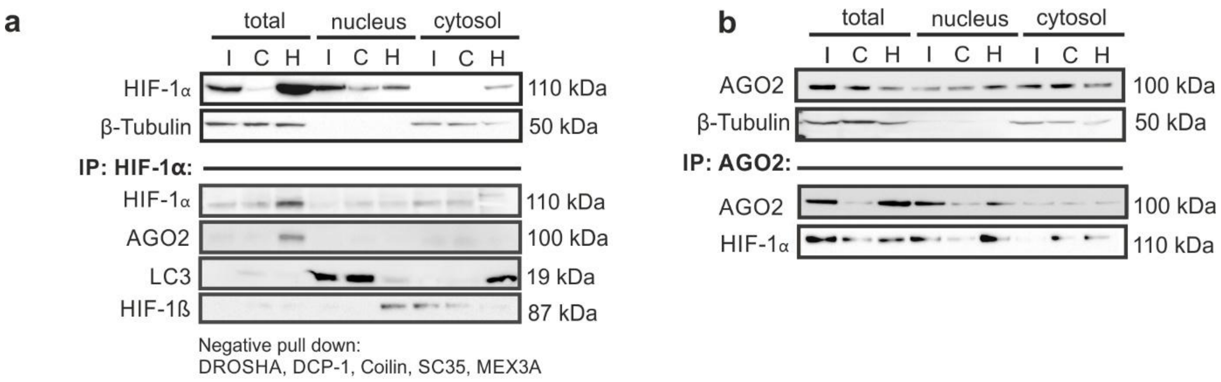


**Supplementary Figure 4. Subcellular analysis of HIF-1α and AGO2 protein interactions**

(a) Immunoblot analysis of HIF-1α-associated protein complexes from total, nuclear, and cytosolic fractions under indicated conditions. Keratinocytes were cultured under normoxic conditions or exposed to hypoxia (1% $O_2$, 24 h). Immunoprecipitation (IP) was performed using anti-HIF-1α antibodies followed by immunoblot analysis for AGO2, LC3, HIF-1β and p300. β-Tubulin served as fractionation control. Negative pull-down controls are indicated.

(b) Immunoblot analysis of AGO2-associated protein complexes from total, nuclear, and cytosolic fractions under indicated conditions. Keratinocytes were cultured under normoxic conditions or exposed to hypoxia (1% $O_2$, 24 h). Immunoprecipitation (IP) was performed using anti-AGO2 antibodies followed by immunoblot analysis for HIF-1α and Dicer. β-Tubulin served as fractionation control.

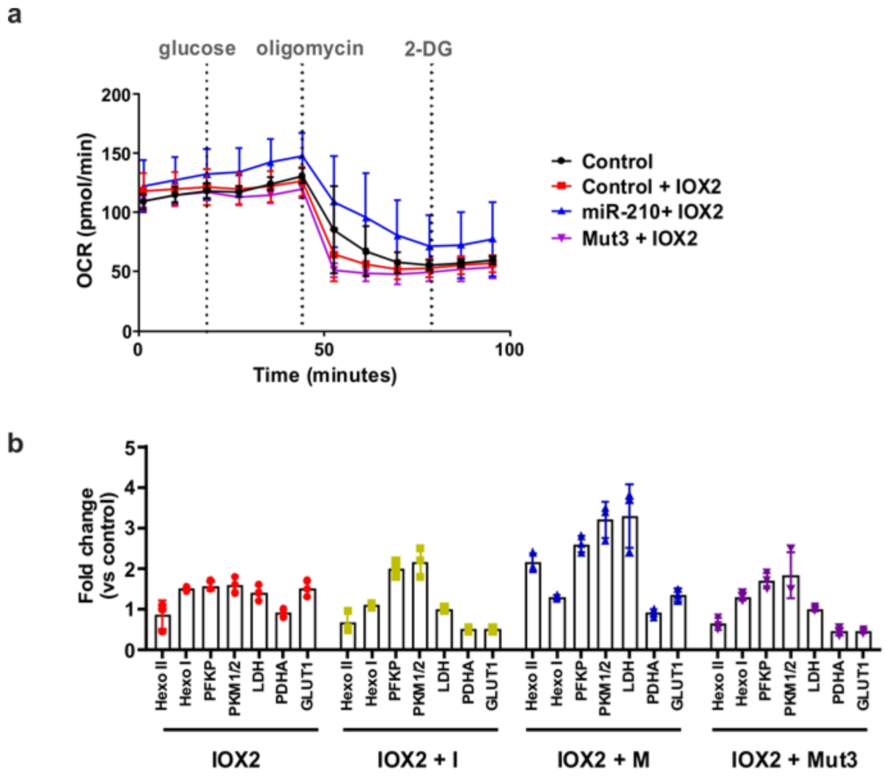


**Supplementary Figure 5. miR-210-5p promotes glycolytic reprogramming without affecting mitochondrial respiration.**

(a) Oxygen consumption rate (OCR) traces of keratinocytes expressing miR-210-5p or the binding-deficient mutant (Mut3) following IOX2 treatment (50 μM, 2 h). Glucose, oligomycin, and 2-deoxy-D-glucose (2-DG) were injected at the indicated time points. Data represent mean ± s.e.m.

(b) Summary of densitometric quantification of glycolytic proteins shown in Figure 5c. Protein expression was normalized to β-tubulin and expressed as fold change relative to untreated control cells (set to 1). Abbreviations: I, miR-210-5p inhibitor; M, miR-210-5p mimic; Mut3, binding-deficient miR-210-5p mutant.

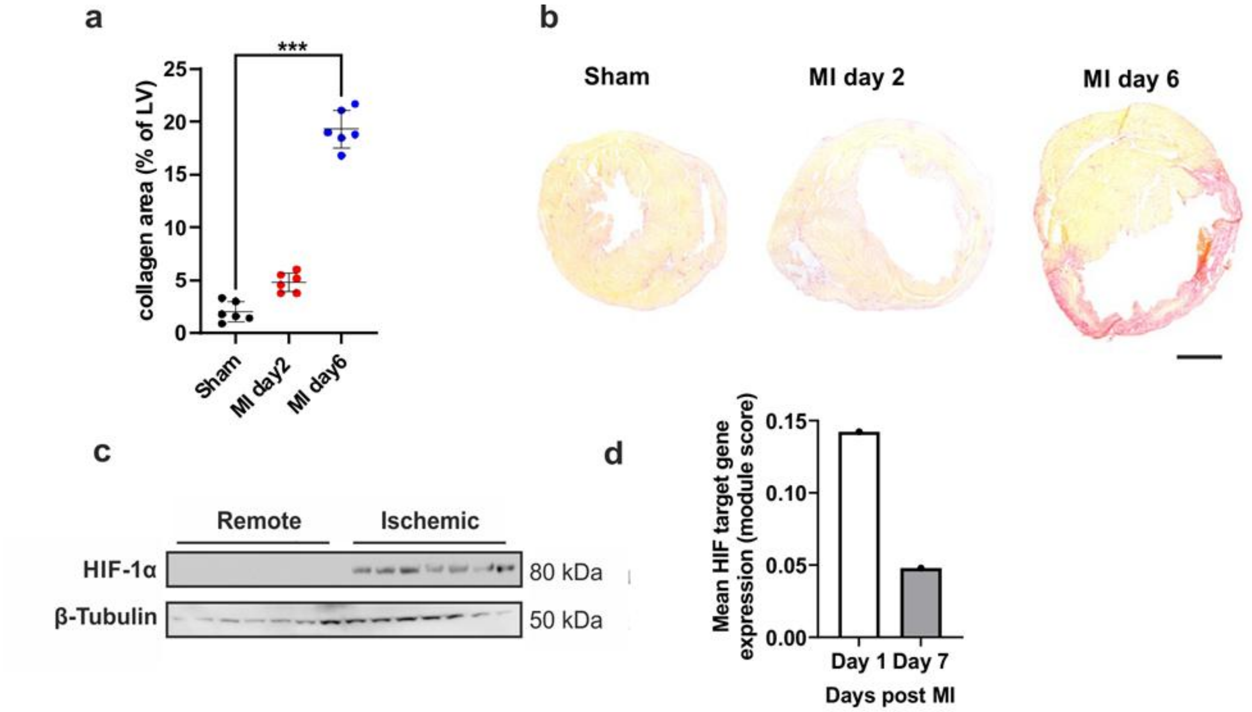


**Supplementary Figure 6. Temporal activation of HIF signaling following murine myocardial infarction.**

(a) Quantification of collagen deposition (percentage of left ventricular area) in sham-operated mice and mice 2 and 6 days after myocardial infarction (MI). Each dot represents one mouse. Statistical analysis was performed using the Kruskal–Wallis test followed by Dunn's multiple-comparisons test. *** p = 0.0003.

(b) Representative histological sections of murine hearts from sham operated controls and at 2- and 6-days post-MI, illustrating progressive collagen accumulation. Scale bar, 1 mm.

(c) Immunoblot analysis of HIF-1α protein levels from remote and ischemic myocardium. β-Tubulin serves as loading control.

(d) Mean expression of canonical HIF target genes in single-nucleus RNA-seq datasets from murine myocardium (GSE214611). Nuclei from sham controls and from hearts collected 1 day (inflammatory phase) and 7 days (reparative phase) post-MI were analyzed. A per-nucleus HIF transcriptional activity score was calculated and averaged per condition.

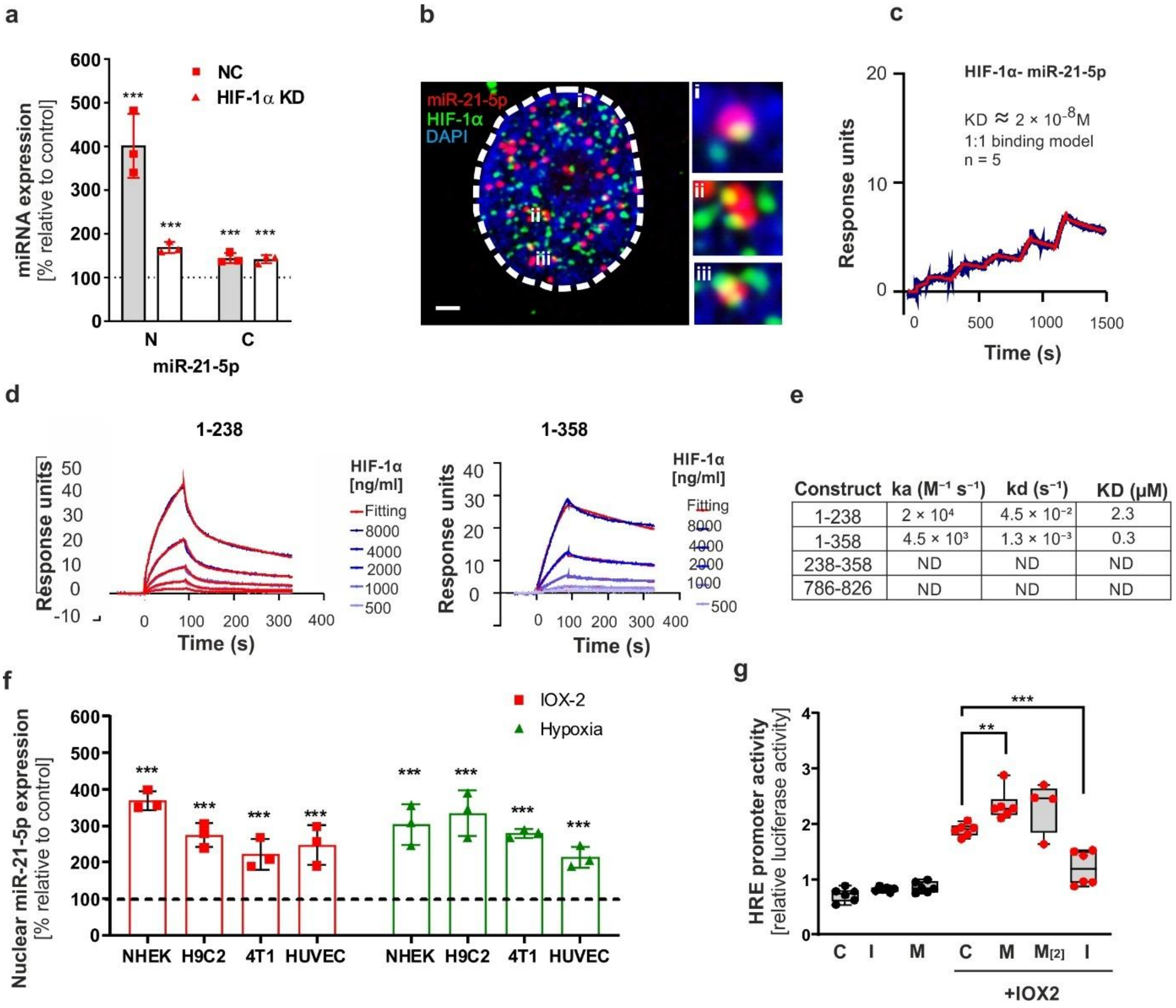


| Construct | ka ($M^{-1} s^{-1}$) | kd ($s^{-1}$) | KD (μM) |
|---|---|---|---|
| 1-238 | $2 \times 10^4$ | $4.5 \times 10^{-2}$ | 2.3 |
| 1-358 | $4.5 \times 10^3$ | $1.3 \times 10^{-3}$ | 0.3 |
| 238-358 | ND | ND | ND |
| 786-826 | ND | ND | ND |

**Supplementary Figure 7. miR-21-5p displays hypoxia-associated nuclear enrichment and directly interacts with HIF-1α.**

(a) Subcellular distribution of miR-21-5p following HIF-1α depletion. Primary human keratinocytes were transfected with control siRNA (NC) or HIF-1α-targeting siRNA (HIF-1α KD), followed by IOX-2 treatment (50 μM, 2 h). Nuclear (N) and cytoplasmic (C) RNA fractions were isolated and analyzed by qPCR for miR-21-5p abundance. Expression levels were normalized to U6 and are shown relative to untreated controls.

(b) Nuclear localization and spatial association of miR-21-5p with HIF-1α. Representative confocal microscopy images showing miR-21-5p detected by RNA-FISH (red) and HIF-1α detected by immunofluorescence staining (green) in keratinocytes exposed to hypoxia (1% $O_2$, 24 h). Insets show enlarged regions highlighting representative nuclear interaction sites. Nuclei were counterstained with DAPI. Scale bars, 5 μm.

(c) Surface plasmon resonance analysis of HIF-1α interaction with miR-21-5p. SPR sensorgrams showing binding of recombinant HIF-1α to immobilized miR-21-5p. Representative sensorgrams and corresponding fitted curves based on a 1:1 binding model are shown.

(d) SPR analysis of HIF-1α truncation constructs interacting with miR-21-5p. Sensorgrams showing interaction of miR-21-5p with recombinant HIF-1α fragments corresponding to amino acids 1–238 and 1–358.

(e) Summary of the association rate constants (ka), dissociation rate constants (kd), and equilibrium dissociation constants (KD) derived from SPR analyses of recombinant HIF-1α truncation constructs interacting with miR-21-5p. ND, no detectable binding.

(f) Nuclear accumulation of miR-21-5p under pharmacological and physiological hypoxia across multiple cell types. NHEK, H9C2, 4T1, and HUVEC cells were treated with IOX-2 (50 μM, 2 h) or exposed to hypoxia (1% $O_2$, 24 h), followed by nuclear RNA isolation and qPCR analysis of miR-21-5p abundance.

(g) HRE reporter activity following modulation of miR-21-5p levels under pharmacological HIF stabilization. Keratinocytes were co-transfected with an HRE-driven firefly luciferase reporter together with control oligonucleotides (C), miR-21-5p mimics (M= 20 nM; M [2] = 40 nM), or miR-210-5p inhibitor (I = 40 nM), followed by IOX-2 treatment (50 μM, 2 h).

Data are presented as mean ± s.e.m. from independent biological replicates. Statistical significance was determined using unpaired two-tailed Student's t-test or one-way ANOVA with appropriate multiple-comparisons correction as described in Methods.

**Supplemental table. 1. Sequence of miRNA inhibitor and siRNA.**

| Gene | Supplier | Sequence (5’) |
|---|---|---|
| *miR-210-5p* Inhibitor | Qiagen | AGTGTGCGGTGGGCAG |
| *Hif1a* siRNA | Dharmacon | - UUUAAUACCCUCCGAUUUA<br>- UUACUGAGUUGAUGGGUUA<br>- GGAAAGAGAGUCAUAGAAC<br>- UGAGAGAAAUGCUUACACA |

**Supplemental table. 2.** List of primers used for quantitative real-time PCR.

| Gene | Forward primer (5’) | Reverse primer (5’) |
|---|---|---|
| *Adipoq* | GGAGAGAAAGGAGATGCAGGT | CTTTCCTGCCAGGGGTTC |
| *Cd36* | TTGTACCTATACTGTGGCTAAATGAGA | CTTGTGTTTTGAACATTTCTGCTT |
| *Egln3* | GTTTGGCTCCCTACCTTGTT | GGATGTCTGCAGGTGTTTCT |
| *Elovl3* | TTCTCACGCGGGTTAAAAATGG | GAGCAACAGATAGACGACCAC |
| *Fabp4* | GGATGGAAAGTCGACCACAA | TGGAAGTCACGCCTTTCATA |
| *Pparg* | TCGCTGATGCACTGCCTATG | GAGAGGTCCACAGAGCTGATT |
| *Ppargc1a* | TTCATCTGAGTATGGAGTCGCT | GGGGGTGAAACCACTTTTGTAA |
| *Serpine1* | TCTGGGAAAGGGTTCACTTTACC | GACACGCCATAGGGAGAGAAG |
| *Tbp* | AGAACAATCCAGACTAGCAGCA | GGGAACTTCACATCACAGCTC |
| *Ucp1* | AGGCTTCCAGTACCATTAGGT | CTGAGTGAGGCAAAGCTGATTT |
| *Vegfa* | AAAAACGAAAGCGCAAGAAA | TTTCTCCGCTCTGAACAAGG |
| *miR-210-3p* | hsa-miR-210-3p (YP00204333)<br>Qiagen cat. Number: 339306 | |
| *miR-210-5p* | mmu-miR-210-5p (YP02105697)<br>Qiagen cat. Number: 339306 | |
| miR-126-5p | Qiagen miScript cat. no. MS00006636 | |
| let-7f | Qiagen miScript cat. no. MS00008274 | |
| *Sno202* | SNORD68 (mmu) (YP00203911)<br>Qiagen cat. Number: 339306 | |

**Supplemental Table 3. List of antibodies.**

| Antigen | Supplier | Catalog |
|---|---|---|
| β-Tubulin | Cell Signaling Technology | 2146 |
| HIF-1α | Cell Signaling Technology | 36169 |
| AGO2 | Abcam | ab57113 |
| LC3 | Cell Signaling Technology | 4599 |
| HIF-1β | Cell Signaling Technology | 5537 |
| p300 | Cell Signaling Technology | 8637 |
| DROSHA | Cell Signaling Technology | 3364 |
| DCP-1 | Abcam | ab47811 |
| Coilin | Cell Signaling Technology | 14168 |
| MEX3A | Abcam | ab79046 |
| Anti-Mouse IgG | Cell Signaling Technology | 7076S |
| Anti-Rabbit IgG | Cell Signaling Technology | 7074S |

**Appendix: Full-length Western Blots**

Full-length, uncropped western blot images for all blots presented in the main figures are provided in the following pages. The cropped regions used in the figures are indicated in blue.

#### Full Blots Figure 2

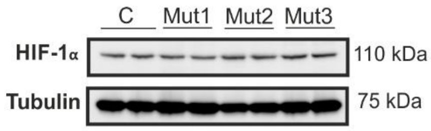


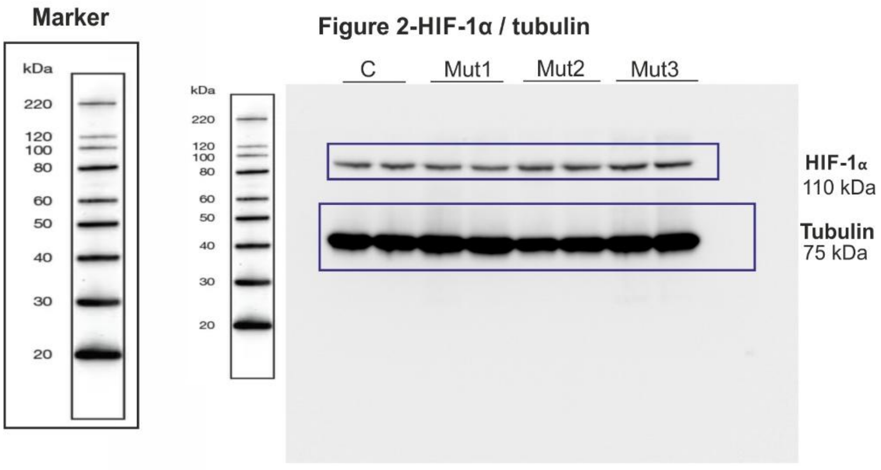

#### Full Blots Figure 5

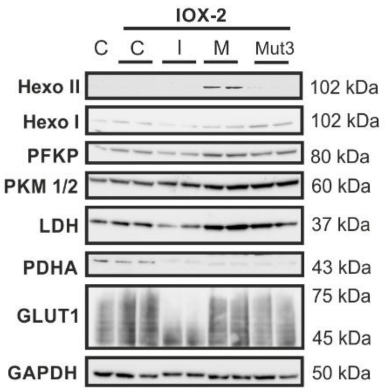


**Marker**

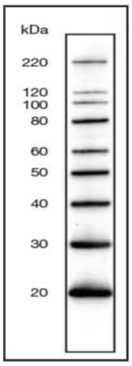


**Figure 5-Hexo I / PFKP / PKM 1/2 / LDH**

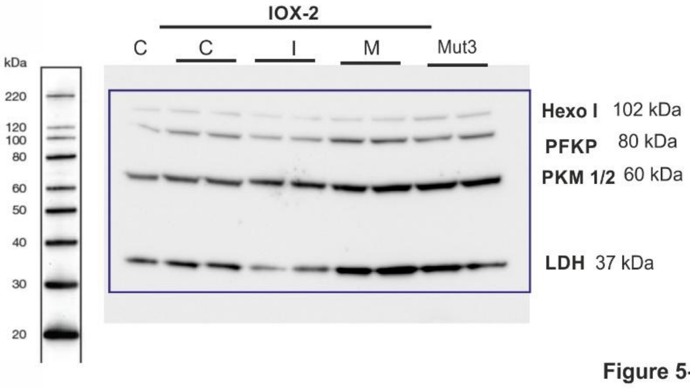


**Figure 5-GAPDH**

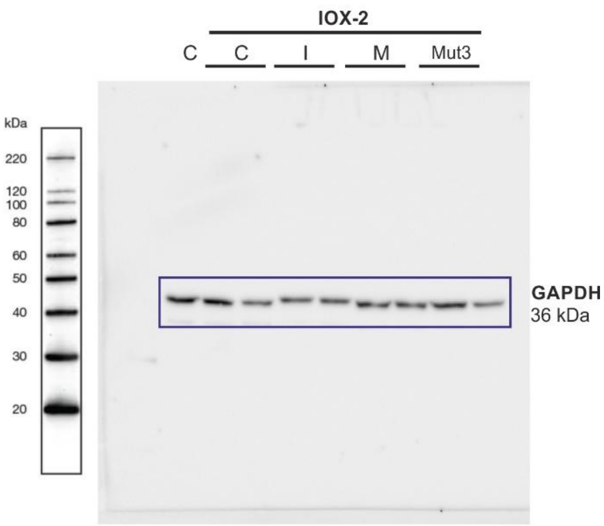


**Figure 5-GLUT1**

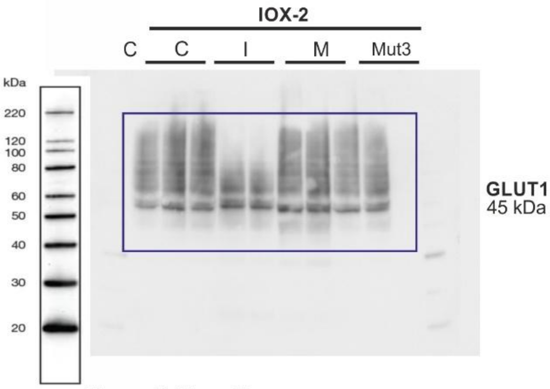


**Figure 5-Hexo II**

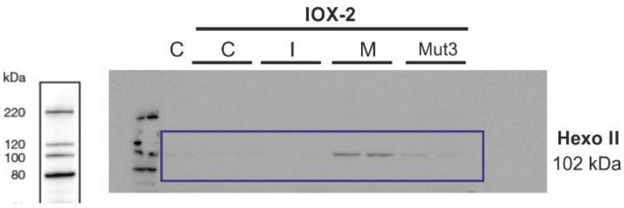

#### Full Blots Supplementary Figure 1

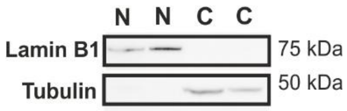


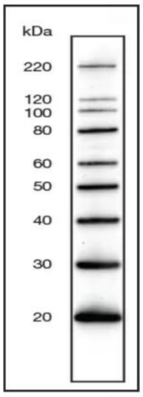


Supplementary Fig. 1 lamin B1 / tubulin

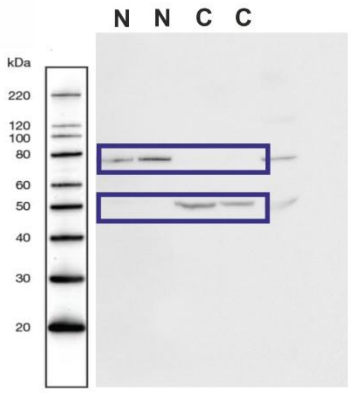

#### Full Blots Supplementary Figure 3A

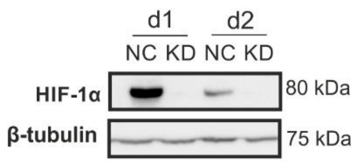


Supplementary Fig. 3A HIF-1α

Marker

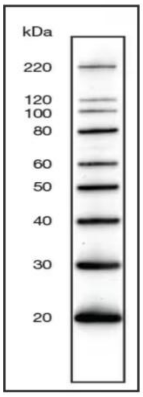


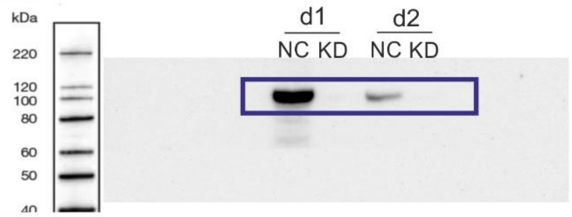


Supplementary Fig. 3A tubulin

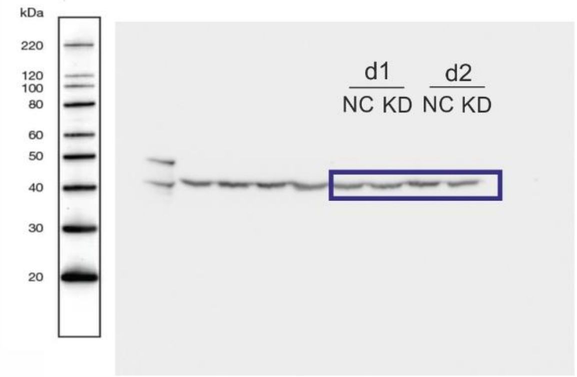

### Full Blots Supplementary Figure 3B

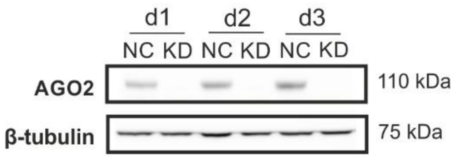


Marker

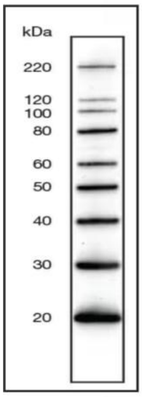


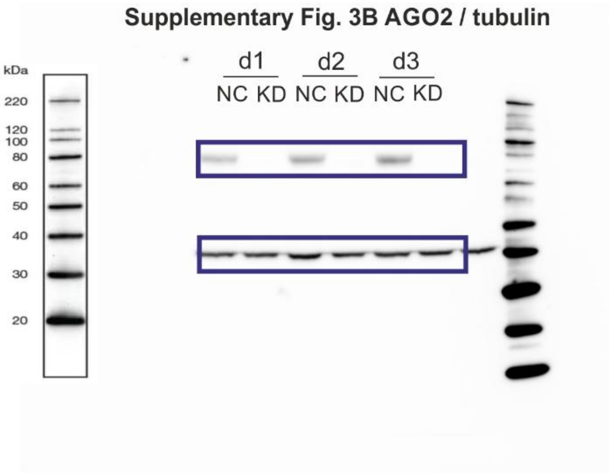

#### Full Blots Supplementary Figure 4a

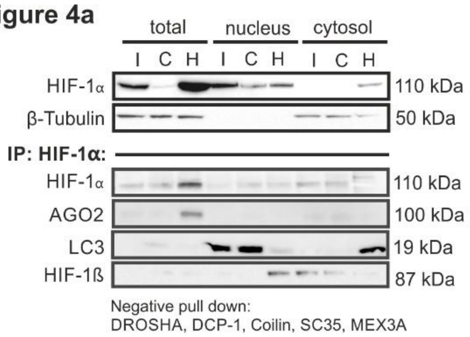


**Marker**

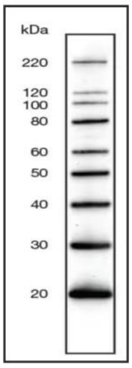


**Figure 4a-HIF-1α / tubulin**

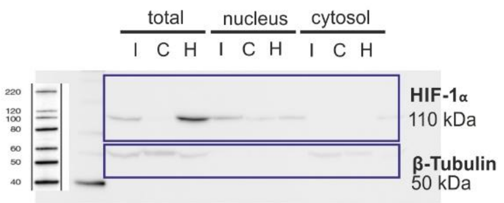


**Figure 4a-HIF-1α**

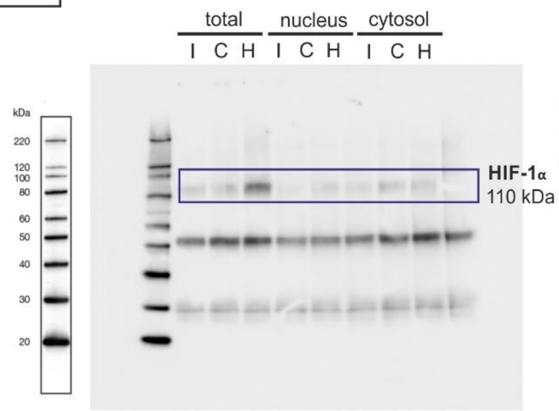


**Figure 4a-HIF-1β**

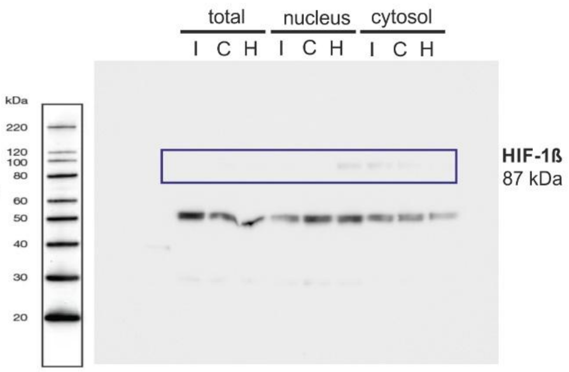


**Figure 4a-AGO-2**

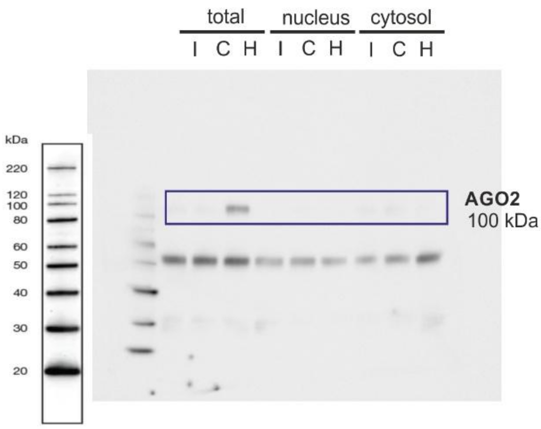


**Figure 4a-LC3**

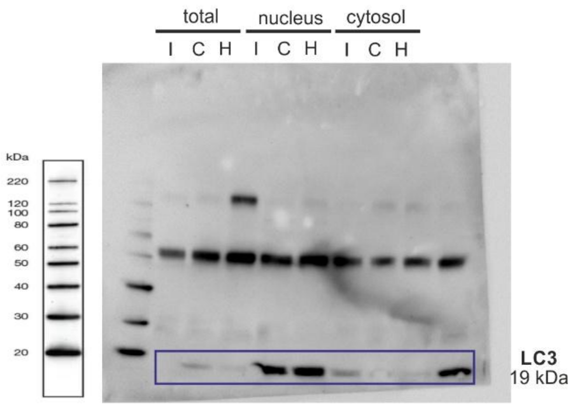

#### Full Blots Supplementary Figure 4b

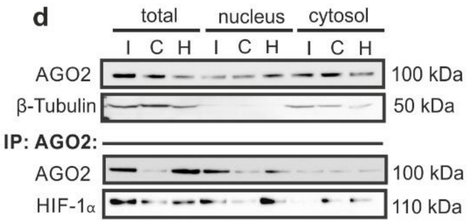


Marker

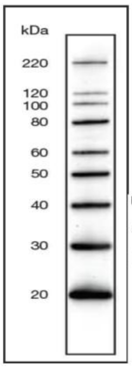


Figure 4b-β-Tubulin

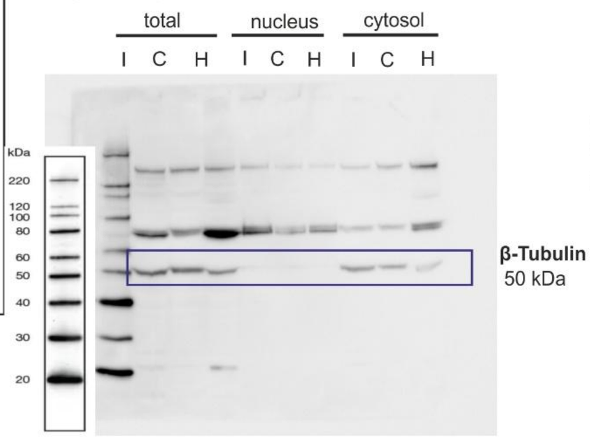


Figure 4b-AGO2

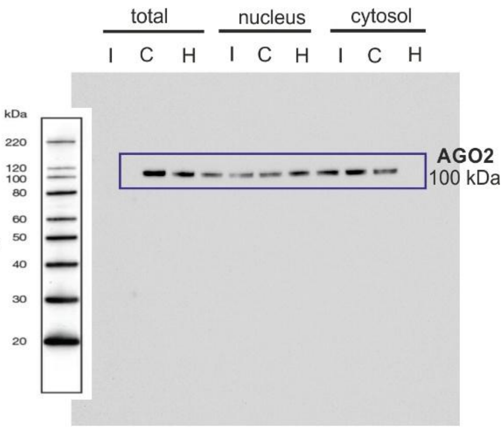


Figure 4b-AGO2

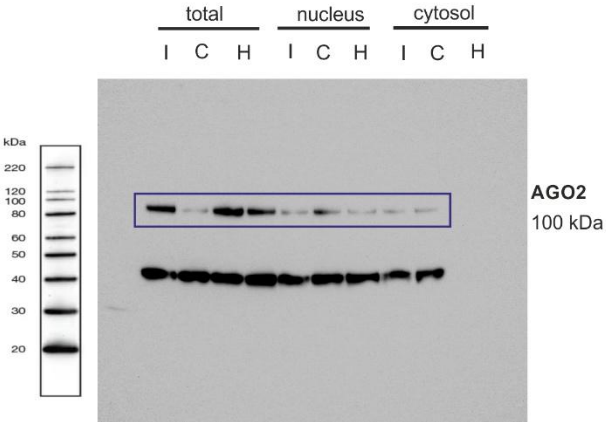


Figure 4b-HIF-1α

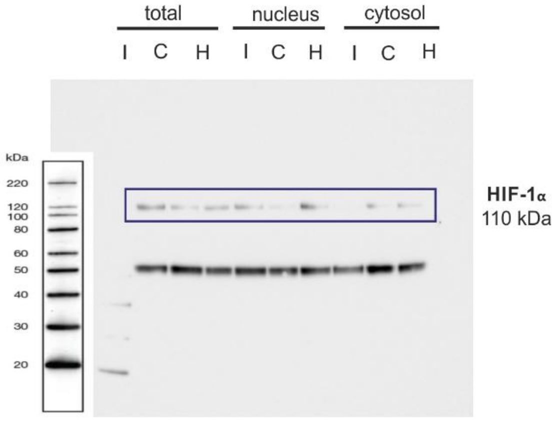

#### Full Blots Supplementary Figure 6

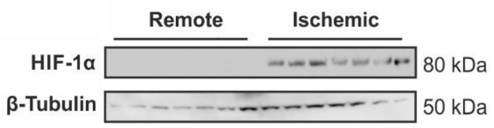


Marker

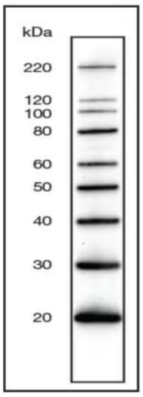


Supplementary Fig. 5 HIF-1α / tubulin

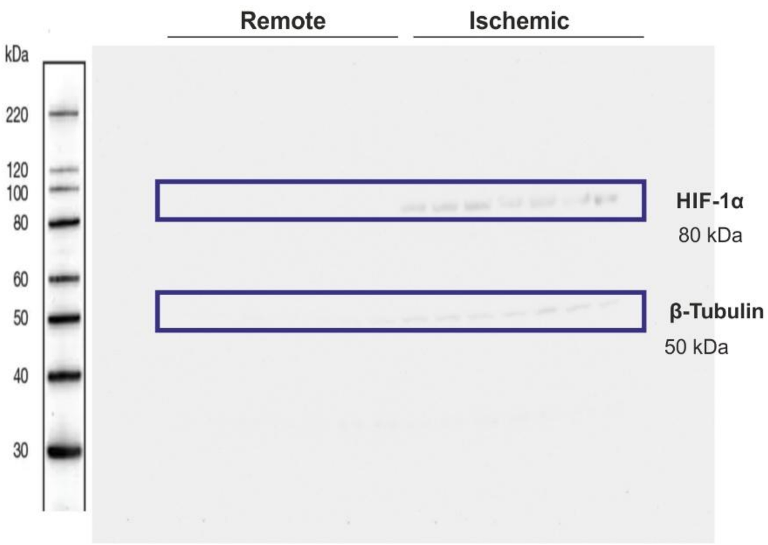
